# RTEL-1 and DNA polymerase theta promote subtelomeric DNA synthesis and telomere fusion in *C. elegans*

**DOI:** 10.1101/2022.09.04.506531

**Authors:** Evan H. Lister-Shimauchi, Morgan Brady, Stephen Frenk, Braxton Harris, Ana-Maria Leon Ortiz, Aylin Memili, Minh Nguyen, Simon Boulton, Shawn Ahmed

## Abstract

Interstitial telomere sequences (ITS) are degenerate telomere tracts scattered along chromosome arms whose functions are not well understood. We found that critically shortened telomeres of *C. elegans* telomerase mutants initiate DNA synthesis within ITS tracts that were close to or far from a telomere. Some ITS tracts were targeted recurrently. RTEL-1 dismantles T-loops and recombination intermediates, and DNA polymerase theta (POLQ-1) promotes end-joining using short segments of microhomology. In telomerase mutants, RTEL-1 and POLQ-1 promoted telomere fusion and DNA synthesis at subtelomeric ITS tracts. RTEL-1 is known to suppress homologous recombination, and we found that RTEL-1 similarly suppressed POLQ-1-mediated double-strand break repair. Mutation signatures characteristic of repair by POLQ-1 occurred during initiation of subtelomeric DNA synthesis and at subsequent template shifting events. We propose that RTEL-1 and POLQ-1 play distinct essential roles in subtelomeric DNA synthesis, a process that may contribute significantly to telomere fusion and tumor genome evolution.

## Introduction

Telomeres are specialized nucleoprotein complexes that cap the ends of linear chromosomes. Telomeres prevent the loss of terminal genome sequences that would otherwise occur because canonical DNA polymerases cannot completely replicate linear chromosomes (Shay & Wright, 2019). Telomeres also prevent chromosome termini from being recognized and repaired as aberrant forms of DNA damage (de Lange, 2018; Smogorzewska & de Lange, 2004).

Telomeres in most metazoans are composed of thousands of base pairs of simple repetitive sequences, such as (TTAGGG)_n_ in mammals and (TTAGGC)_n_ in *C. elegans*. Telomerase maintains telomere length by reverse transcribing *de novo* telomere repeats onto chromosome termini (Collins, 2006). Telomerase is present at high levels in germ cells and is expressed in some human somatic stem cells. However, most human somatic cells display little or no telomerase activity due to silencing of the TElomerase Reverse Transcriptase (TERT) locus (Shay & Wright, 2019). This results in a ‘telomere clock’ that limits the number of divisions that human somatic cells can undergo before entering senescence in response to DNA damage induced at shortened telomeres (Chiba et al., 2015; Heidenreich et al., 2014; Horn et al., 2013). Silencing of somatic telomerase is a tumor suppressor mechanism that was created on multiple occasions during the evolution of large mammals, in order to control the proliferation of vast numbers of somatic cells (Gomes et al., 2011; Seluanov et al., 2007). TERT promoter mutations are among the most prominent recurrent mutations in cancer, which illustrates the importance of telomere-induced senescence as a tumor suppressor mechanism (Chiba et al., 2015; Heidenreich et al., 2014; Horn et al., 2013).

Inactivation of checkpoint genes that promote senescence leads to further cell proliferation, ultimately resulting in critically shorted telomeres that induce pronounced genomic instability and high levels of cell death, termed ‘crisis’ (Cleal et al., 2018; Maciejowski & de Lange, 2017). Critically shortened telomeres (‘uncapped telomeres’) of human somatic cells that bypass senescence can fuse to create ‘chromatid-type fusions’ if sister chromatids fuse during G2 or ‘chromosome-type fusions’ if telomeres of distinct chromosomes fuse during G1 (Cleal & Baird, 2020). The resulting dicentric chromosomes can be pulled to opposite sides of a spindle during mitosis, which can trigger breakage-fusion-bridge cycles (McClintock, 1941). Cells that emerge from crisis can maintain their telomeres by expressing TERT, which activates telomerase, or by activating the telomere maintenance pathway termed Alternative Lengthening of Telomeres (ALT) (Pickett & Reddel, 2015).

Genome rearrangements that occur in response to uncapped telomeres are likely to be prominent features of developing tumors (Cleal et al., 2018). Mechanisms relevant to creation of end-to-end fusions can be studied in mammalian cells (Maciejowski et al., 2015), but analysis of genome rearrangements that occur at human telomeres has been complicated by the repetitive nature of many sub-telomeres (Baird, 2018; Linardopoulou et al., 2005). In contrast, the nematode *C. elegans* has only 6 chromosomes whose subtelomeres are largely unique (Lowden et al., 2008). In addition, *C. elegans* chromosomes are holocentric, which allows for end-to-end chromosome fusions that occur in strains that are deficient for the *C. elegans* telomerase reverse transcriptase *trt-1* to be outcrossed and maintained as stable genome rearrangements in the presence of telomerase (Meier et al., 2006). End-to-end chromosome fusions created in *trt-1* mutants can display subtelomeric duplications that are either simple duplications or composites of template-shifting events (Lowden et al., 2011). Therefore, a promiscuous DNA synthesis process commonly adds segments of subtelomeric DNA from a sister chromatid to critically shortened telomeres prior to the creation of chromosome-type end-to-end fusions (Lowden et al., 2011).

ITSs are mixtures of perfect and imperfect telomere repeat sequences that are common features of metazoan chromosome arms (Lin & Yan, 2008; Meyne et al., 1990). Critically shortened telomeres of *C. elegans* telomerase mutants sometimes initiate DNA synthesis at a subtelomeric ITS tract prior to the creation of a telomere fusion (Lowden et al., 2011). Template shifting within a subtelomere can occur during DNA synthesis events at ITS tracts (E. Kim et al., 2021; Lowden et al., 2011). Similar template switching events have been observed during meiosis in *S. cerevisiae (Lee et al., 2014; Stafa et al., 2014; Tsaponina & Haber, 2014)* and contribute to mitotic and meiotic mammalian genome rearrangements (Hartlerode et al., 2016; Koumbaris et al., 2011; Zhang et al., 2009).

Telomeres end in 3’ single-stranded overhangs that can create strand-invasion intermediates by folding back and interacting with a segment of double-stranded telomeric DNA (Griffith et al., 1999; Tomáška et al., 2020). The resulting T-loop structures protect chromosome termini from being recognized as DNA damage and can be stabilized by Shelterin components such as the double-stranded telomere binding protein TRF2 in mammalian cells and by the single-stranded telomere binding protein POT-1 in *C. elegans* (de Lange, 2018; Raices et al., 2008; Van Ly et al., 2018). The RTEL1 helicase promotes telomere stability via anti-recombinase functions that unwind T-loops in the context of telomere replication (Fig. 5A) and that dismantle stable four-stranded G-quadruplex DNA structures and associated R-loops that can form during telomere replication (Kotsantis et al., 2020; Vannier et al., 2012). RTEL1 also represses homologous recombination during meiosis by disrupting D-loops (Barber et al., 2008; León-Ortiz et al., 2018; Vannier et al., 2014; Youds et al., 2010). Biallelic RTEL1 mutations occur in humans that suffer from Hoyeraal-Hreidarsson syndrome, which is a strong single-generation disorder that presents early in life and reflects a pronounced deficiency for telomere maintenance (Ballew, Joseph, et al., 2013; Ballew, Yeager, et al., 2013; Deng et al., 2013; Guen et al., 2013). In addition, heterozygous RTEL1 mutations can induce telomere instability in germ cells, resulting in transgenerational telomere erosion in human families that is reminiscent of what occurs when humans are haploinsufficient for reverse transcriptase or RNA subunits of telomerase (Kannengiesser et al., 2015).

DNA polymerase theta (POLQ in mammals or POLQ-1 in *C. elegans*) utilizes microhomology to repair DNA double-strand breaks (Chan et al., 2010; van Kregten & Tijsterman, 2014; Wood & Doublié, 2016). *C. elegans* POLQ-1 has been shown to create small genomic deletions in the absence of the DOG-1 helicase, where one breakpoint is adjacent to a long tract of guanine-rich DNA that has been deleted (Koole et al., 2014). These small deletions are thought to occur when the replisome encounters an impassable DNA structure and represent strong evidence for the existence and toxicity of G-quadruplex structures *in vivo*. Short tracts of homology are typically found at genomic deletion breakpoints created by POLQ-1 - typically 2 to 20 nucleotides in mammals and a single nucleotide in *C. elegans* (Black et al., 2016; Kent et al., 2016; Koole et al., 2014; Van Schendel et al., 2015; Wyatt et al., 2016). POLQ promotes biogenesis of chromosome-type end-to-end fusions in response to acute telomere uncapping, where long telomeres that lack TRF2 are sealed as head-to-head tracts of long double-stranded telomeric DNA with short patches of microhomology (Mateos-Gomez et al., 2015).

Here we study the biogenesis of chromosome fusions that are created in response to critically shortened telomeres of *C. elegans* telomerase mutants. Our results suggest that RTEL-1 and POLQ-1 play prominent roles in telomere-ITS DNA synthesis events, some of which are recurrent, and in the creation of telomere fusions.

## Results

### Telomeres target ITSs adjacent to a subset of *C. elegans* chromosome termini

Previous analysis of end-to-end chromosome fusions isolated from *C. elegans* mutants deficient for telomerase indicated that critically shortened telomeres can initiate DNA synthesis at ITS tracts that are 8 to 50 kilobases from a telomere (Lowden et al., 2011). We decided to empirically study telomere-ITS DNA synthesis events using Polymerase Chain Reaction (PCR) assays that detect fusion of a telomere with an ITS tract. Both ends of Chromosome *IV* were excluded from our assays because highly repetitive sequences immediately adjacent to each telomere preclude PCR amplification (Lowden et al., 2008). We therefore studied 5 ITS tracts that were closest to each telomere of Chromosomes *I*, *II*, *III*, *V,* and *X* (Fig. 1A). We developed primer pairs that robustly amplified all 50 subtelomeric ITSs. We used primers facing the right telomere of Chromosome *X* (*XR*) and the left telomere of Chromosome *V* (*VL*) to amplify previously characterized telomere-ITS DNA synthesis events present in *ypT29* and *ypT50* strains that possess *XR-IL* and *XR-VL* end-to-end chromosome fusions, respectively (Lowden et al., 2011). Primers facing the *XR* telomere and the first ITS tract of *IL or VL* (oriented towards the 3’ end of the G-rich strand of the ITS tract, Fig. S1A) amplified the telomere-ITS DNA synthesis events present in genomic DNA from *ypT29* and *ypT50* strains, even when genomic DNA containing these chromosome fusions was diluted 1:100 with N2 wild type genomic DNA. No PCR products were observed for template genomic DNA from wild type or from an early-generation *trt-1* telomerase mutant strain that lacked chromosome fusions (Fig. S1B-C). PCR is therefore a sensitive means of detecting telomere-ITS DNA synthesis events.

**Figure 1.**
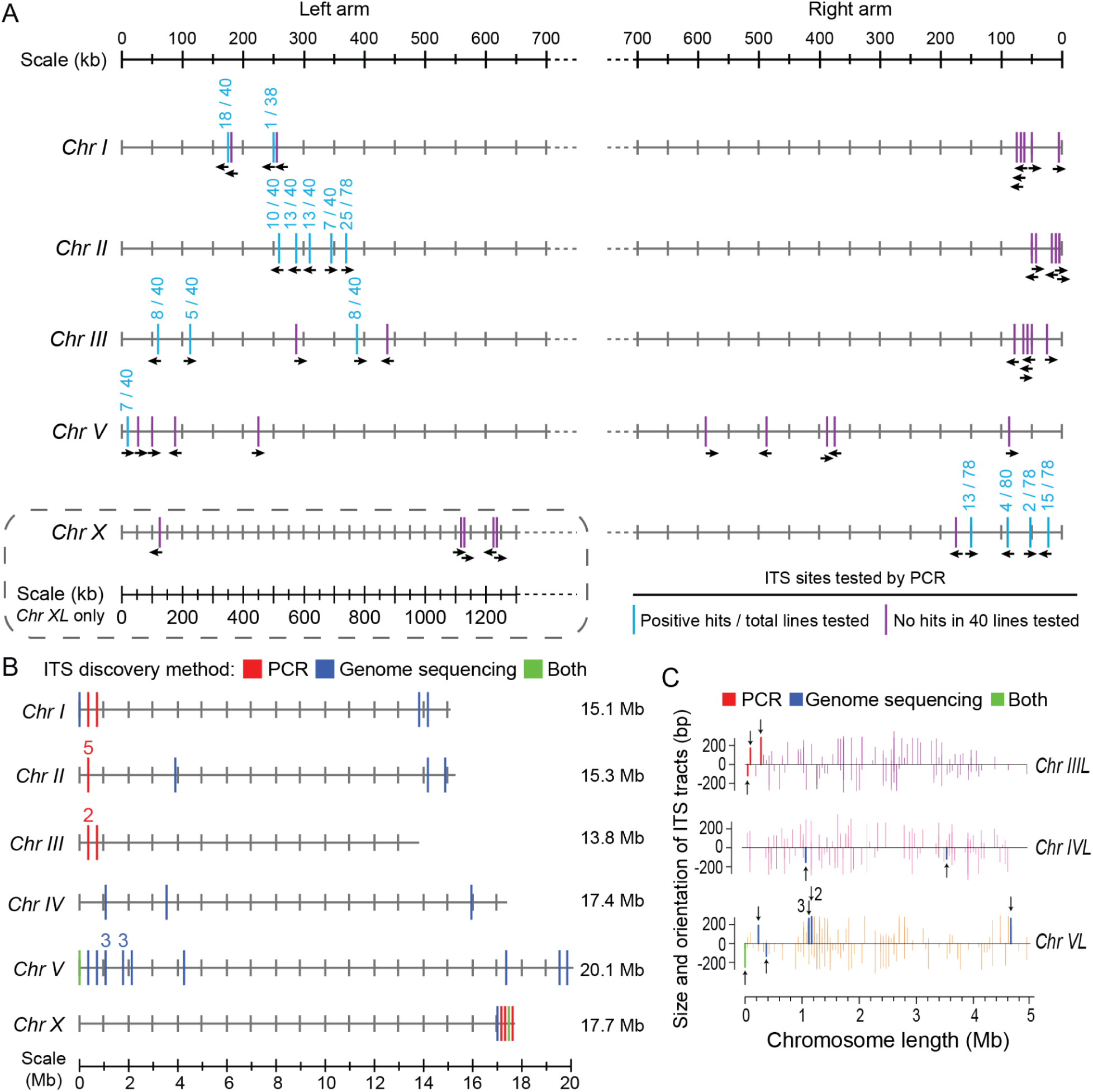
Analysis of telomere-ITS recombination events in genomes of *C. elegans trt-1* telomerase mutants. (A) PCR assays of 5 terminal ITS tracts at 10 *C. elegans* telomeres reveal ITS tracts that can be targeted by critically shortened telomeres (blue), as well as ITS tracts that were tested using validated primers but not detected as being targeted (purple). Numbers above ITS tracts indicate the number of recombination events detected over the total number of lines tested. The five closest ITS tracts to each chromosome end are shown, except for Chromosome *IL*, where one tract very close to the telomere was censored from PCR. ITS tract orientation with respect to each telomere is shown by black arrows. (B) Genome sequence analysis reveals many additional ITS tracts that can be targeted by shortened telomeres. Blue ITS tracts were identified only by genome sequence analysis, red ITS tracts were identified only by PCR, and green ITS tracts were identified by both techniques. (C) Telomere-ITS recombination events at the left ends of Chromosomes *III*, *IV* and *V* are shown as thick lines surrounded by thin ITS tracts that were resistant to recombination with shortened telomeres. Numbers refer to independent telomere-ITS recombination events that occurred at either the same ITS tract or were too close together to show as distinct lines.

We thawed a *trt-1* telomerase mutant strain that was created by outcrossing the *trt-1* mutation twenty times in a heterozygous condition using an *unc-29* balancer mutation, which possesses telomeres that are initially wild-type in length (Meier, 2006). We grew 40 independent lines of this *trt-1* mutant strain in parallel and harvested genomic DNA when roughly 30 to 40 progeny were produced by 6 parents grown on a single plate, which indicates that the brood size has dropped substantially due to the creation of multiple end-to-end chromosome fusions (Ahmed & Hodgkin, 2000; Meier et al., 2006). By comparison, 6 wild-type worms will produce about one thousand progeny. We performed PCR with the 50 telomere-ITS primer pairs that target the 5 terminal ITS tracts of Chromosomes *I*, *II*, *III*, *V,* and *X*, and found that we could detect PCR products for ITS tracts at a single end of each *C. elegans* chromosome tested (Fig. 1A). We confirmed these results with 40 independently grown *trt-1* mutant lines and observed similar frequencies of telomere-ITS DNA synthesis (Fig. S1D-E). The ability of an ITS tract to undergo telomere-ITS DNA synthesis did not depend on its orientation with respect to the telomere (Fig. 1A).

To obtain an unbiased perspective on the fate of critically shortened telomeres, we performed Illumina short-read whole genome sequencing using genomic DNA isolated from late-generation *trt-1* mutant lines and from *trt-1* mutant lines that had activated the telomerase-independent telomere maintenance pathway ALT. These strains all contained multiple end-to-end chromosome fusions (Cheng et al., 2012; Meier et al., 2006). We defined telomere-ITS DNA synthesis events in these genomes based on duplication or amplification of reads adjacent to the longest stretch of perfect telomere sequence within an ITS tract (Fig. 2) (Lowden et al., 2011). The detected DNA synthesis events targeted ITS tracts that were not distributed randomly across all chromosome arms (*P*=0.027, Shapiro-Wilks test for normality). The right arm of *Chromosome III* and the left arm of *Chromosome X* may be resistant to telomere-ITS DNA synthesis (Fig. 1B). Four of 21 novel ITS sites were found >2 MB from a telomere (Fig. 1C).

**Figure 2.**
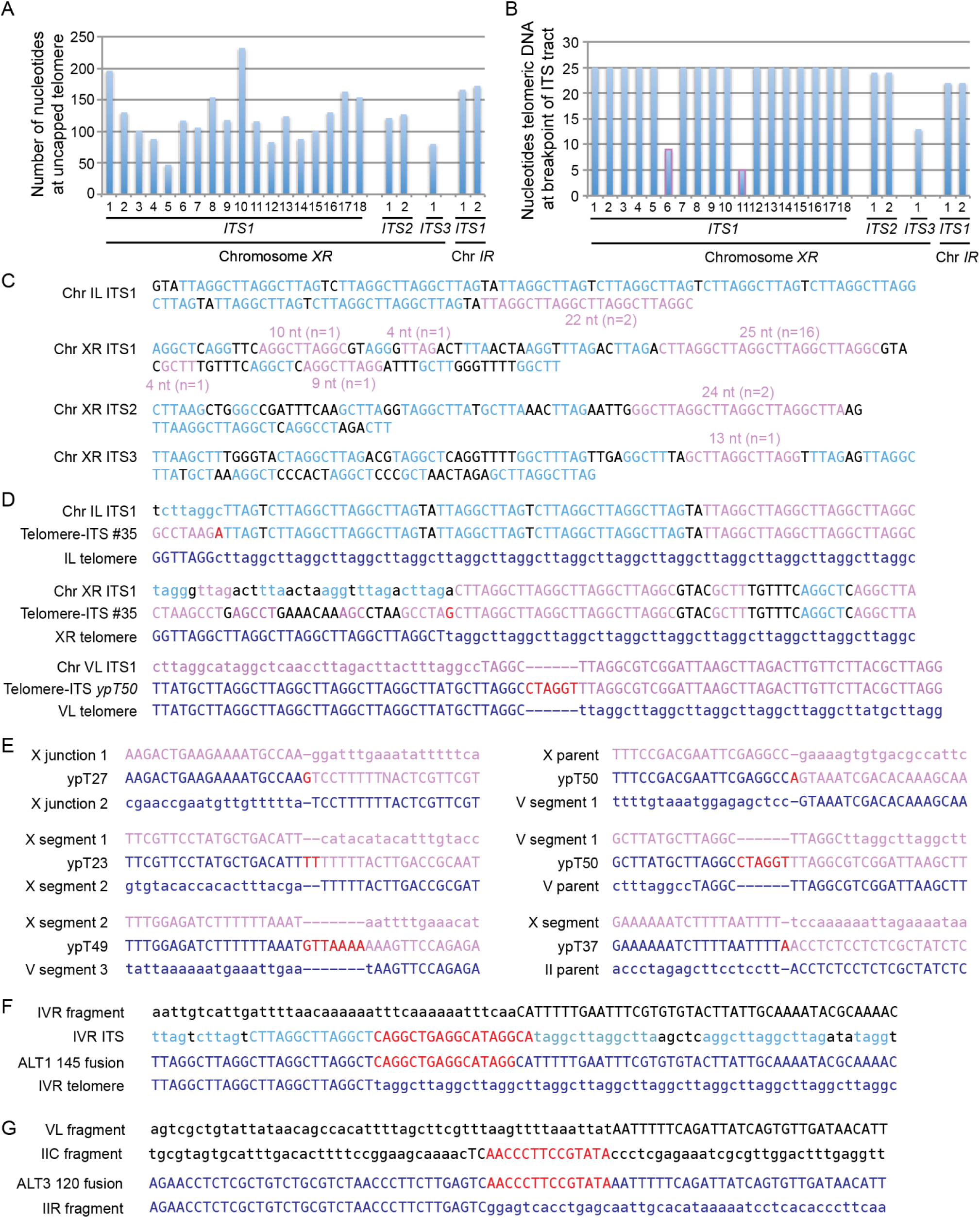
Critically shortened telomeres target long tracts of telomere homology. (A) Lengths of critically shortened telomeres at the time of telomere-ITS fusion. (B) Length of perfect telomere sequences within ITS tracts that are targeted by critically shortened telomeres. Most events concern the longest tract of perfect telomere homology within an ITS tract (blue), whereas several events target shorter tracts (lavender outline). (C) Telomere-ITS DNA synthesis events begin at lavender telomere sequences within four ITS tracts. Light blue text depicts > 2 nt homology to TTAGGC telomere sequence within ITS tracts. (D) Telomere-ITS DNA synthesis events with single nucleotide changes (D) or insertions (E) at transitions between telomere and ITS sequences shown in red. Sequence of original telomere shown in dark blue. Sequence present at telomere-ITS DNA synthesis events shown in capital letters. Upper and lower sequences represent original DNA sequences prior to DNA synthesis, lower case letters are missing after telomere-ITS DNA synthesis and dashes represent sites where de novo nucleotide insertion occurs. (F) ALT1 145 contig reveals that a IVR-IIIL telomere fusion fragment began when a IVR telomere initiated DNA synthesis at an ITS tract (red sequence within IVR ITS) and then shifts to black sequence at a distinct subtelomeric location (IVR fragment). (G) Template-shifting event during creation of IIR-VL telomere fusion that begins at IIR fragment locus (ALT3 120 contig), then transitions to the red sequence copied from the IIC fragment locus, then transitions to VL subtelomere (VL fragment).

### Uncapped telomeres initiate DNA synthesis at sites of perfect telomere homology

We sequenced 23 telomere-ITS DNA synthesis breakpoints at 4 distinct ITS tracts (Fig. 2C), primarily sampling events at the first ITS tract on the right end of the *X* chromosome. We calculated the length of perfect telomere sequence present at the time of telomere-ITS DNA synthesis based on the total number of perfect telomere repeats where DNA synthesis began, which assumes maximal annealing of the 3’ overhang of a dysfunctional telomere with a stretch of perfect telomere sequence in an ITS tract. Critically shortened telomeres possessed 47 to 232 bp of telomere repeats, with an average length of 127 nucleotides (Fig. 2A). Consistently, the maximal amount of telomeric DNA identified in telomere fusions from human fibroblasts was 252 base pairs, with a mean length of 35 base pairs (Letsolo et al., 2010). Independent studies found 48 to 430 bp of telomeric DNA at sites of telomere fusion in human papillary neoplasms or 100 to 800 bp of telomeric DNA from human breast cancer samples (Hata et al., 2018; Tanaka et al., 2012). These results are generally consistent with minimal lengths of double-stranded telomeric DNA that promote T-loop formation *in vitro* and *in vivo* (Griffith et al., 1999; Stansel et al., 2001). Functional human telomeres can be as short as 300 bp in length (Allen & Baird, 2009; Britt-Compton et al., 2006, 2009), which is only slightly longer than the minimal length necessary for telomere stability in *C. elegans*.

Most telomere-ITS PCR products identified from late-generation *trt-1* mutant genomic DNA revealed that DNA synthesis initiated at the longest stretches of perfect telomere sequence within an ITS tract, which were 25, 24, 22 and 13 nucleotides in length (Fig. 2B-C). However, a few ITS DNA synthesis events initiated at shorter segments of telomere homology that were 4, 4, 9 and 10 nucleotides in length (Fig. 2C and S2A). Telomere-ITS DNA synthesis events #10 and #35 displayed single nucleotide mutations at transitions between telomere and ITS DNA sequences (Fig. 2D). In addition, a previously described telomere-ITS DNA synthesis event, *ypT50* (Lowden et al. 2011), displayed a 6 nt insertion at the transition between telomere and ITS sequences (Fig. 2D). These mutations suggested the promiscuous DNA synthesis activity of DNA polymerase theta (Ramsden & Carvajal-Garcia, 2021).

Telomere fusions from *trt-1* telomerase mutants were previously shown to display subtelomeric template shifting events (Lowden et al., 2011). We found that roughly half of these template shifting events possessed small insertions at transitions between segments of subtelomeric DNA (Fig. 2E, red text). Several telomere fusion events in the genomes of *C. elegans trt-1* telomerase mutants that survive via ALT were recently reported based on long-read Nanopore sequencing (E. Kim et al., 2021). We observed that the *IVR* telomere of the ALT1 145 telomere fusion initiated DNA synthesis at a nearby ITS tract (red sequence) and that a template-shifting event then occurred at a distinct nearby sequence that possessed 2 nucleotides of homology (Fig. 2F). A distinct template-shifting occurred at the *IIR-VL* telomere fusion breakpoint (ALT3 120 contig), where a IIR subtelomeric fragment initiated DNA synthesis at a short segment of DNA from the center of chromosome *II* (*IIC*) and then transitioned to the *VL* subtelomere (Fig. 2G). Deletions or translocations that are bridged by a short patch of DNA that is copied from elsewhere in the genome are characteristic of DNA polymerase theta, and so are short 1 or 2 nucleotide segments of homology present at these template-shifting events (Koole et al., 2014; Van Schendel et al., 2015; van Schendel et al., 2016). Consistently, we observed similar short segments of homology in previously described subtelomeric template-shifting events that occurred during creation of *C. elegans* telomere fusions (Fig. S2B) (Lowden 2011).

### The ERCC-1 nuclease does not alter telomere-ITS DNA synthesis

Double-minute chromosomes are extrachromosomal DNA circles that result in oncogene amplification in tumors (Murnane, 2006). Deficiency for the ERCC1 endonuclease subunit in telomerase-positive mouse cells promotes biogenesis of double-minute chromosomes, possibly because ERCC1 cleaves strand invasion intermediates that occur between telomeric 3’ overhangs and ITS tracts (Zhu et al., 2003). In addition, loss of ERCC1 in the context of telomerase deficiency in *Arabidopsis* leads to accelerated sterility, possibly because telomeric termini recombine with ITS tracts (Vannier et al., 2009). In both of these examples, telomere-ITS DNA synthesis events were hypothesized to occur in the absence of ERCC1, but these events were not verified by DNA sequence analysis. To ask if loss of ERCC-1 affects DNA synthesis events at *C. elegans* ITS tracts, we created *trt-1 ercc-1* double mutant strains utilizing the *ercc-1* deletions *tm1981* and *tm2013*. We propagated 40 strains of each *trt-1 ercc-1* double mutant and observed that sterility was substantially accelerated in comparison to *trt-1* single mutants (Fig. S3A), consistent with *Arabidopsis* telomerase mutants that are deficient for ERCC1 (Vannier et al., 2009). When *trt-1 ercc-1* double mutant brood size had dropped substantially, we prepared genomic DNA from each strain and carried out PCR assays for all ITS tracts that were targeted for DNA synthesis by short telomeres (Fig. 1A). We observed that telomere-ITS DNA synthesis occurred at a frequency similar to that observed for late-generation *trt-1* single mutants (Fig. 3A). Previous suggestions that ERCC1 might repress telomere-ITS DNA synthesis in mammals and plants could reflect a distinct role of ERCC1 in modulating telomere stability (Vannier et al., 2009; Zhu et al., 2003).

**Figure 3.**
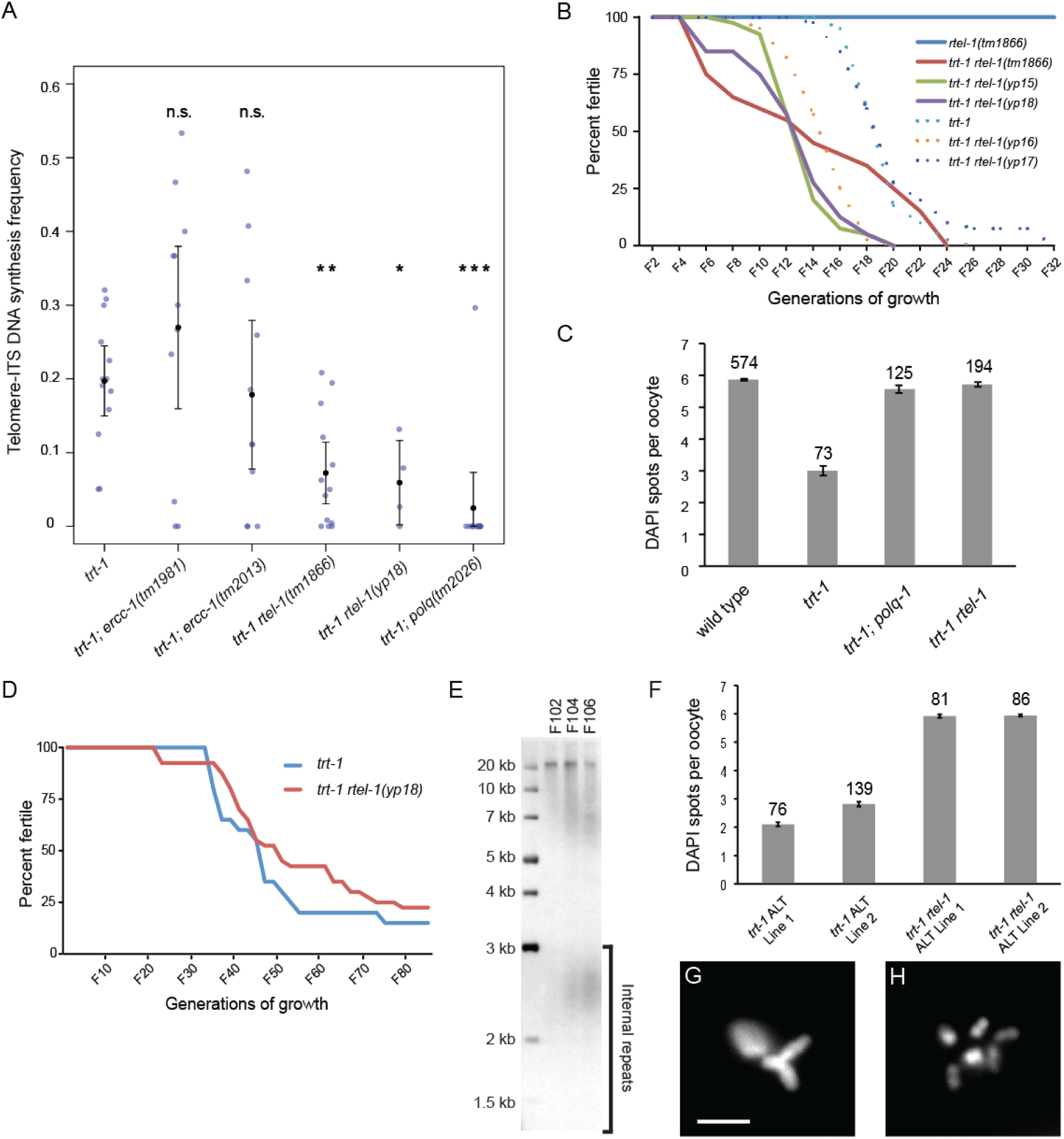
RTEL-1 and POLQ-1 promote telomere-ITS DNA synthesis and end-to-end chromosome fusion. (A) Telomere-ITS DNA synthesis frequency is strongly reduced by deficiency for *rtel-1* and *polq-1* but is unaffected by deficiency for *ercc-1.* Each point represents one chromosome arm for a single sample set. Significance represents comparison with *trt-1* single mutants. (B) Loss of *rtel-1* shortens the time to sterility in a *trt-1* mutant background. Dotted lines represent *trt-1* mutant backgrounds with at least partial *rtel-1* function, as the *yp16* and *yp17* alleles are not predicted to be loss-of-function alleles. (C) High levels of end-to-end chromosome fusions were observed in *trt-1* mutants but were absent in late generation *trt-1 rtel-1* and *trt-1; polq-1* double mutants. Significance represents comparison with wild type. (D) RTEL-1 is not required for *trt-1* telomerase mutants to survive via the ALT telomere maintenance pathway, which can be induced when large numbers of worms are transferred. (E) Southern blotting reveals telomere maintenance in *trt-1; rtel-1* double mutant strains that survive via ALT. (F) RTEL-1 promotes formation of end-to-end chromosome fusions in the context of ALT. (G, H) Representative images showing reduced chromosome numbers in *trt-1* ALT lines (G) and wild-type chromosome numbers in *trt-1 rtel-1* ALT lines (H). Scale bar represents 5 µm. * p < 0.05, ** p < 0.01, *** p < 0.001.

### POLQ-1 and RTEL-1 promote telomere-ITS recombination and end-to-end chromosome fusion

Disruption of telomere binding proteins can lead to fusion of long mammalian telomeres via a pathway termed alternative-non-homologous end-joining (a-NHEJ) or theta-mediated end-joining (TMEJ) (Badie et al., 2015; Kratz & de Lange, 2018; Mateos-Gomez et al., 2015; Rai et al., 2010, 2019; Sfeir & de Lange, 2012). A-NHEJ/TMEJ relies on DNA polymerase theta to heal a DNA double-strand break by annealing a 3’ overhang with a short 1-6 nucleotide stretch of microhomology, typically at a nearby target sequence but sometimes from a sequence that is distant or even on another chromosome (Chan et al., 2010; van Kregten & Tijsterman, 2014; Wood & Doublié, 2016). One characteristic of DNA polymerase theta action is the addition of small insertions at breakpoints that it heals, typically 1-25 nucleotides that are usually copied from a nearby DNA sequence (Koole et al., 2014; Van Schendel et al., 2015). Several breakpoints of subtelomeric DNA synthesis events documented this study and in genome sequences we examined from an independent study of *C. elegans* telomerase mutants displayed insertions and sequence transitions that are hallmarks of DNA polymerase theta (Fig. 2D-G and S2) (E. Kim et al., 2021; Lowden et al., 2011). We therefore created *trt-1; polq-1* double mutants, passaged 27 lines until their fertility had dropped substantially (Fig. S3), prepared genomic DNA, and used PCR to test ITS tracts that we previously observed were targeted by short telomeres (Fig. 1A). We found that telomere-ITS DNA synthesis was markedly reduced in the absence of *polq-1* (Fig. 3A).

Loss of RTEL1 in mouse cells results in creation of telomere circles by the structure-specific nuclease SLX4 because RTEL-1 fails to dismantle T-loops (Vannier et al., 2012). Moreover, RTEL-1 suppresses homologous recombination, likely by dismantling strand-invasion events that resemble T-loops (Barber et al., 2008; León-Ortiz et al., 2018; Youds et al., 2010). We therefore hypothesized that RTEL-1 might suppress telomere-ITS DNA synthesis events by dismantling telomere-ITS strand invasion intermediates. We tested this hypothesis by creating *trt-1 rtel-1(tm1866)* double mutants, which displayed the low levels of fertility previously reported for *rtel-1(tm1866)* single mutants (Fig. S4) (Barber et al., 2008). We confirmed this result by creating independent CRISPR/Cas9-induced *rtel-1* mutations in a *trt-1* mutant background (Arribere et al., 2014; Dickinson et al., 2013)*. rtel-1* deletion mutations *yp15* and *yp18* are expected to cause frameshifts in *rtel-1* transcripts, both of which resulted in significantly reduced fertility, comparable to that of *rtel-1(tm1866)* single mutants (Fig. S4). Although deficiency for telomerase suppresses accumulation of telomeric damage in RTEL1-deficient mouse cells (Margalef et al., 2018), we found that deficiency for telomerase did not affect the fertility defect of *C. elegans rtel-1* mutants. The reduced fertility of *C. elegans rtel-1* single mutants may instead reflect a distinct function of RTEL-1 in antagonizing homologous recombination or facilitating global DNA replication (Kotsantis et al., 2020; León-Ortiz et al., 2018; Vannier et al., 2013; Youds et al., 2010).

We propagated *trt-1 rtel-1(tm1866)* double mutants and observed that the onset of sterility was faster than for *trt-1* single mutants, although the difference was not statistically significant (Fig. 3B, Holm adjusted p = 0.1174). We tested this potential genetic interaction with independent CRISPR/Cas9-induced *rtel-1* deletion mutations *yp15* and *yp18*, both of which displayed a significantly faster onset of sterility in the absence of *trt-1* (Fig. 3B, Holm adjusted p < 10^-12^ for both). These results indicate that RTEL-1 promotes telomere stability in the absence of telomerase, which is consistent with the rapid sterility observed for *Arabidopsis* RTEL1 mutants that are deficient for telomerase (Recker et al., 2014). We tested a role for RTEL-1 telomere-ITS DNA synthesis events by examining all 50 ITS tracts sampled in Fig. 1A and found that genomic DNA from late-generation *trt-1 rtel-1* double mutants displayed markedly diminished levels of telomere-ITS DNA synthesis (Fig. 3A) (*trt-1* vs *trt-1 rtel-1(tm1866)*: p = 0.0021, n = 120 and 116 lines; *trt-1* vs *trt-1 rtel-1(yp18)*: p = 0.020, n = 120 and 38 lines).

### RTEL-1 and POLQ-1 are essential for biogenesis of telomere fusions

Given that loss of *rtel-1* and *polq-1* compromised telomere-ITS DNA synthesis, we asked if these mutations perturbed the creation of telomere fusions that are normally observed in late-generation telomerase mutants. We counted DAPI-stained chromosome bodies in oocytes in late-generation *trt-1 rtel-1* and *trt-1; polq-1* double mutant strains. We found that *trt-1* late-generation single mutant controls typically displayed 3 or 4 DAPI bodies per oocyte that indicate the presence of end-to-end chromosome fusions, consistent with previous observations (Fig. 3C) (Ahmed and Hodgkin 2000; Meier et al. 2006; Meier et al. 2009). However, late-generation *trt-1 rtel-1* and *trt-1; polq-1* double mutants displayed 5.72+/-0.07 and 5.57+/-0.12 DAPI body counts, which were only modestly decreased in comparison to the 5.86+/-0.03 found in wild-type control oocytes (*trt-1 rtel-1* vs wild type p = 0.0003, *trt-1; polq-1* vs wild type p = 10^-5^). This suggests that telomere fusions rarely occur when *polq-1* or *rtel-1* are deficient (Fig. 3C). Consistently, mouse cells that are deficient for both RTEL1 and telomerase lack telomere fusions, at least for early-passage mouse embryonic fibroblasts (Margalef et al., 2018).

Polymorphisms near the human RTEL1 locus have been previously linked to cancer and occur in the context of ALT-positive tumors (Wrensch et al. 2009; Mirabello et al. 2010), and telomere-ITS DNA synthesis events have been previously implicated in the creation of *C. elegans* telomerase mutant strains that survive via the ALT pathway (E. Kim et al., 2021; Seo et al., 2015). We therefore asked if loss of *rtel-1* in *C. elegans* would affect the induction of ALT, which can be activated if *C. elegans trt-1* telomerase mutants are grown under crowded conditions (Cheng et al., 2012). However, when we transferred crowded *trt-1 rtel-1(yp18)* double mutant strains by chunking agar plates once a week for over 80 generations, the fraction of *trt-1 rtel-1* lines that survived via ALT-mediated telomere maintenance was similar to that of *trt-1* single mutants (Fig. 3D; p=0.373). It remains possible that *C. elegans rtel-1* might affect the induction of ALT in absence of the histone H3.3 chaperone ATRX/DAXX, which is almost always deficient in human ALT tumors (Cheng et al., 2012; Heaphy et al., 2011; Lovejoy et al., 2012; Schwartzentruber et al., 2012).

Southern blotting revealed that *trt-1 rtel-1* ALT strains maintain telomere length in the absence of telomerase (Fig. 3E), as previously observed for *trt-1* single mutant ALT strains (Cheng et al., 2012). As *C. elegans trt-1* telomerase deficient strains that survive typically experience multiple chromosome fusions prior to induction of ALT (Cheng et al., 2012), we examined *trt-1 rtel-1* ALT strains and found that they possessed 5.9+/-0.1 DAPI bodies per oocyte, as observed for wild-type *C. elegans* strains (Fig. 3F-H) (Boerckel et al., 2007; Meier et al., 2006, 2009). Therefore, RTEL-1 is required for biogenesis of telomere fusions created in response to telomerase deficiency, even for ALT strains that have been passaged for many generations. These data indicate that activation of ALT in *C. elegans* germ cells does not depend on the biogenesis of telomere fusions (Cheng et al., 2012).

### DNA polymerase theta heals CRISPR-induced breaks in the absence of RTEL-1

RTEL-1 antagonizes HR and DNA polymerase theta is capable of functionally substituting for HR (Barber et al., 2008; Ceccaldi et al., 2015; León-Ortiz et al., 2018; Mateos-Gomez et al., 2015; Wyatt et al., 2016; Youds et al., 2010). Given that loss of both *polq-1* and *rtel-1* resulted in shared phenotypes that include reduced levels of telomere-ITS DNA synthesis and lack of telomere fusions in telomerase mutants, we asked if RTEL-1 might be required for POLQ-1-mediated TMEJ by inducing CRISPR/Cas9 edits in the gene *dpy-10*. We found that *rtel-1* mutants displayed a significantly higher frequency of CRISPR-induced *dpy-10* mutations in comparison to wild-type controls (Fig. 4A). Sequencing of resulting *dpy-10* mutants (wild type n=15, *rtel-1* n=18) revealed small deletions as well as deletions accompanied by small insertions in both *rtel-1* mutant and wild-type control backgrounds (Fig. 4B and 4C, Table S1). Insertions in both strains that were longer than 7 nucleotides possessed matches to local or distant segments of the genome, indicating that RTEL-1 is not required for the ability of POLQ-1 to initiate DNA synthesis at a distant segment of the genome and then shift back to seal a DSB (Fig. 4D). As the overall frequency of CRISPR-induced mutations is reduced 6-fold when *polq-1* is mutant, and as small deletions accompanied by insertions are eliminated in the absence of POLQ-1 (Van Schendel et al., 2015), we conclude that RTEL-1 is not required for healing of DSBs by POLQ-1.

**Figure 4.**
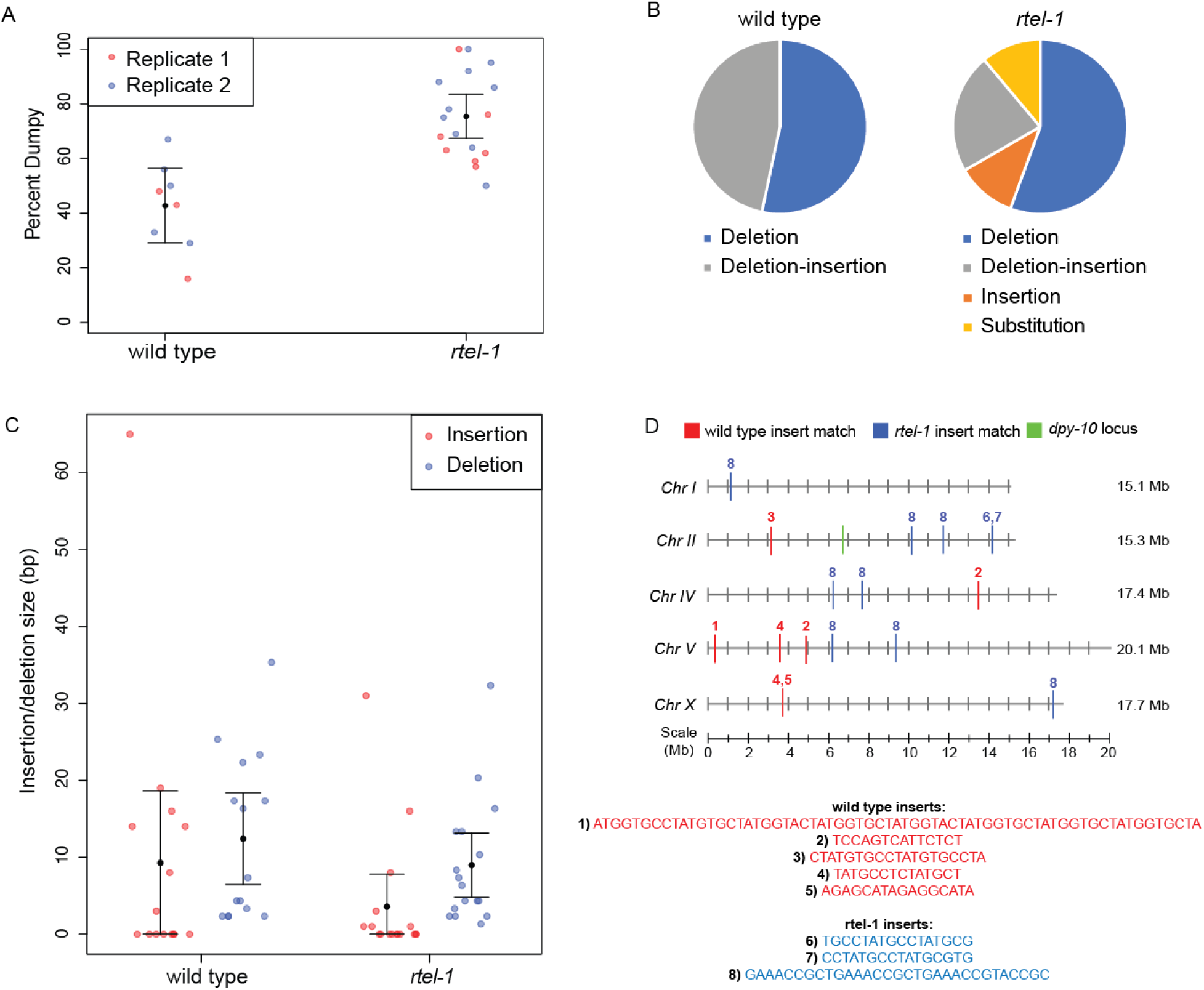
RTEL-1 is not required for Pol-theta-mediated DNA repair. (A) After injection with CRISPR constructs targeting *dpy-10* and an mCherry injection marker, the percentage of mCherry+ F1 worms that displayed a dumpy phenotype was significantly higher in *rtel-1* mutants compared to wild-type controls (Wilcox test, p<0.0005). (B) DNA repair after *dpy-10* targeting in wild-type and *rtel-1* mutant worms yielded both insertions and deletions. (C) Deletion and insertion sizes were not significantly different between N2 and *rtel-1* mutants. (D) Insertions in both wild-type (red) and *rtel-1* mutant (blue) worms aligned to sequences throughout the genome. The *dyp-10* locus is shown in green. Insert sequences are shown below.

## Discussion

We tested fifty ITS tracts close to ten telomeres and found that critically shortened telomeres create ITS DNA synthesis events at five telomeres, typically within 300 Kb of a *C. elegans* telomere (Fig. 1A). Whole genome sequence analysis of genomic DNA of late-generation *trt-1* mutants revealed only 2 out of 23 telomere-ITS DNA synthesis events targeted the fifty ITS tracts closest to ten telomeres (Fig. 1B, Table S2). Instead, the majority of ITS tracts targeted by short telomeres are >1 Mb from a telomere. This suggests that telomeric 3’ overhangs of short uncapped telomeres can scan vast territories of subtelomeric DNA in order to identify a segment of perfect homology to the *C. elegans* telomere repeat (TTAGGC)_n_. Consistently, the length of T-loops is highly variable in organisms with long segments of double-stranded telomeric DNA, which implies that overhangs of normal telomeres possess an inherently flexible mechanism for scanning long stretches of double-stranded telomeric DNA prior to creation of a strand invasion intermediate (Doksani et al., 2013; Griffith et al., 1999; Tomáška et al., 2020; Van Ly et al., 2018).

The orientation of an ITS tract relative to a telomere did not affect its susceptibility to DNA synthesis (Fig. 1A, Fig. 2C). We asked if epigenomic signatures, such as H3K9 di- or tri-methylation or H3K4 trimethylation, were associated with the susceptibility of ITS tracts to telomere-ITS DNA synthesis events but could not find evidence for this (not shown). ITS tracts at most chromosome ends were capable of undergoing telomere-ITS DNA synthesis (Fig. 1B), except for the right end of Chromosome *III* and the left end of Chromosome *X*. The telomere at the left end of the Chromosome *X* is immediately adjacent to a 10 Kb palindrome, and repetitive sequences next to this palindrome are often targeted for DNA synthesis by uncapped *XL* telomeres (Lowden et al., 2011). Subtelomeric sequence composition might therefore suppress telomere-ITS DNA synthesis by channeling uncapped telomeres into alternative telomere repair outcomes.

Short telomeres of proliferating human fibroblasts can induce DNA synthesis at 46 nucleotides of perfect telomere homology within an ITS tract (Capper et al., 2007). However, the relevance of the vast repertoire of ITS tracts to the fate of critically shortened telomeres is not understood. We observed that a subset of terminal ITS tracts are targeted for DNA synthesis by dysfunctional telomeres, often recurrently, which implies that similar recurrent DNA synthesis events may shape genomes of developing tumors. Note that an ITS tract occurs near the promoter of the human TERT gene, which is in close physical proximity to the telomere of Chromosome 5p and is frequently activated during tumor development(Arias-Pulido et al., 2002; Garnis et al., 2005).

Mammalian telomeres have been proposed to influence gene expression by interacting with ITS tracts via loops (W. Kim et al., 2016). Telomere-ITS associations could be mediated by telomeric proteins that physically interact with ITS tracts (Aksenova & Mirkin, 2019). The double-stranded telomere binding protein TRF1 associates with interstitial telomere repeats in hamster cells (Krutilina et al., 2001; Smogorzewska et al., 2000). The human 2q14 ITS tract was created by a head-to-head telomere fusion event (Ijdo et al., 1991) and interacts with TRF1, TRF2, and other shelterin proteins. Aphidicolin or knockdown of TRF1 induce fragility at 2q14 but not at other non-telomeric fragile sites (Bosco & de Lange, 2012). On the other hand, TRF2 and RTEL1 but not TRF1 have been shown to repress DNA damage foci at pericentromeric repeats and other heterochromatic segments of the genome, which may reflect roles for TRF2 and RTEL1 in relieving topological stress that occurs when DNA replication forks encounter a T-loop (Mendez-Bermudez et al., 2018). Roles for telomere binding proteins in responding to replication stress and topological stress at ITS tracts are consistent with the presence of perfect and degenerate telomere repeats within ITS tracts (Bosco & de Lange, 2012; Mendez-Bermudez et al., 2018; Sarek et al., 2016). The sequence and protein composition of ITS tracts may make them attractive targets for DNA synthesis events that are initiated by telomeres that are too short to form a T-loop. If telomerase is not available and a telomere is too short to form a T-loop, then associated telomere binding proteins that normally chaperone T-loop formation may promote intercalation of the 3’ overhang within a subtelomeric ITS tract (Fig. 5B). If this strand-invasion intermediate possesses >7 nucleotides of perfect homology to the telomeric overhang, this might represent a suitable substrate for initiation of DNA synthesis by a replicative polymerase like DNA polymerase delta. However, natural chromosome termini inhibit DNA repair at sites of DNA damage near or within telomeres. Telomeres resemble a one-sided DNA double-strand break and are coated with proteins that inhibit local DNA damage responses including ATM signaling (TRF2), ATR signaling (POT1a), canonical non-homologous end-joining (TRF2), and alternative non-homologous end-joining (shelterin and Ku) (Denchi & de Lange, 2007; Hockemeyer et al., 2006; Mateos-Gomez et al., 2015; Wu et al., 2006). In addition, if senescence is induced in human somatic cells in response to ionizing radiation, irreparable forms of DNA damage near telomeres can persist for months (Fumagalli et al., 2012). Therefore, a strand-invasion intermediate between the 3’ overhang of a short telomere and a stretch of perfect telomere homology within an ITS tract may possess ‘telomeric qualities’ that repress initiation of DNA synthesis by DNA polymerase delta.

**Figure 5.**
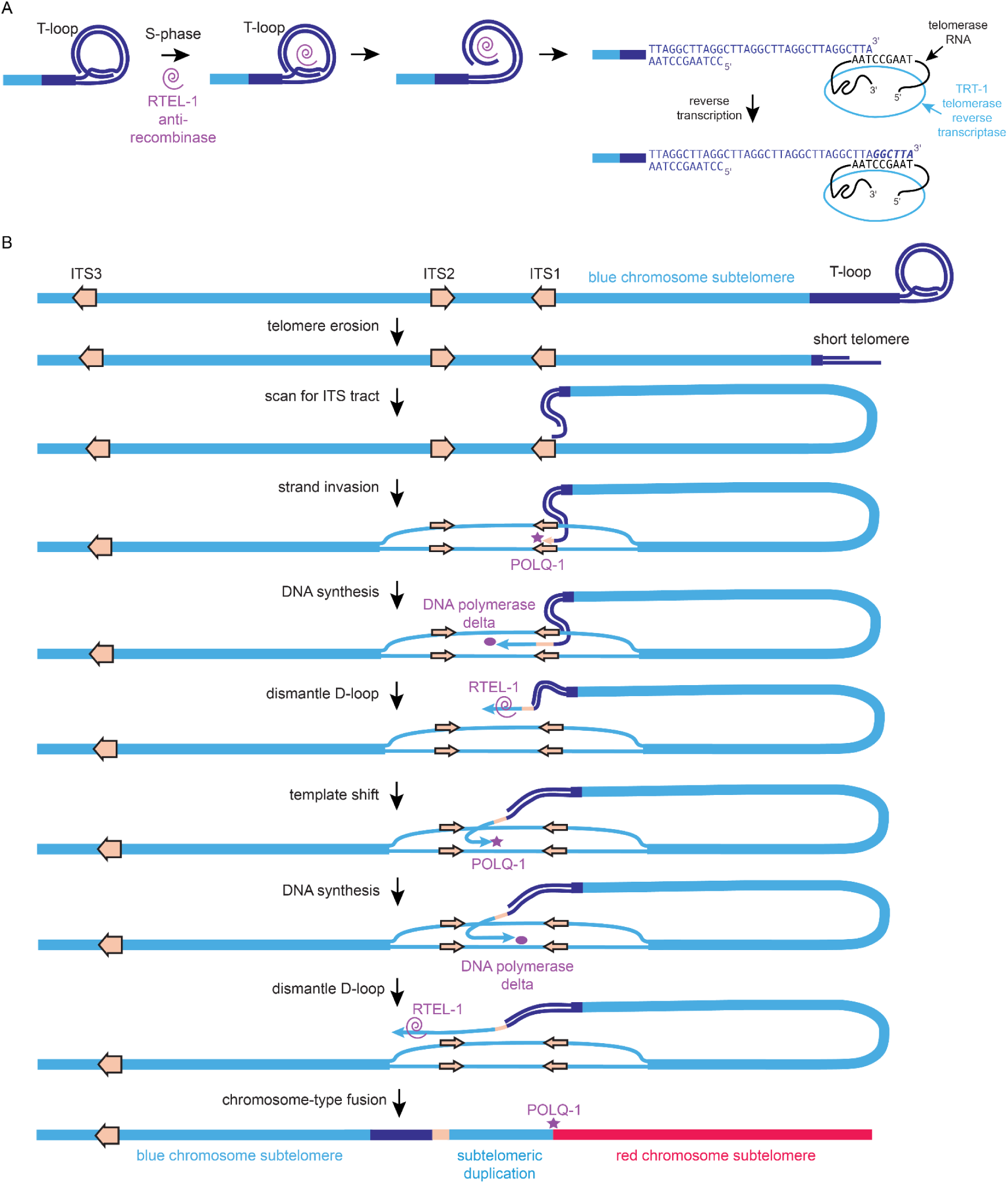
RTEL-1 and POLQ-1 promote telomere-ITS DNA synthesis when telomerase is deficient. (A) RTEL-1 can act as an anti-recombinase that dismantles T-loops when telomeres are replicated, which may provide telomerase access to the chromosome terminus. The terminal nucleotide at telomeric 3’ overhangs and the sequence of the telomerase RNA template are not yet known. (B) When telomeres become critically short in the absence of telomerase, telomeric 3’ overhangs will scan chromosome arms and anneal to the longest stretch of perfect telomere repeat homology within an ITS tract. POLQ-1 initiates subtelomeric DNA synthesis at this site, DNA polymerase delta extends the primer created by POLQ-1, and the strand displacement activity of RTEL-1 promotes a template-shifting event prior to a second round of strand-displacement that allows POLQ-1 to create a chromosome-type fusion with a sub-telomere of a distinct chromosome.

RTEL-1 normally dismantles T-loops as telomeric DNA is replicated during S-phase, which may set the stage of telomere repeat addition by telomerase (Fig. 5A). In the absence of telomerase, a telomere that is too short to form a T-loop may form a strand invasion intermediate at an ITS tract that could be resistant to the activity of DNA polymerase delta (Fig. 5B). In this circumstance, RTEL-1 might dismantle this unusual strand-invasion intermediate, perhaps when it encounters a DNA replication fork, and hand the 3’ overhang to DNA polymerase theta, which might then create a short primer that would allow DNA polymerase delta to initiate subtelomeric DNA synthesis. DNA polymerase theta prefers substrates that possess long 3’ overhangs (Wyatt et al., 2016) and might have a natural affinity for overhangs of critically short telomeres whose stability might improve if subtelomeric DNA synthesis were initiated.

Homopolymeric tracts of guanines in the *C. elegans* genome can form very stable G-quadruplex structures that trigger 50 bp - 200 bp deletions (Cheung et al., 2002), which are created by DNA polymerase theta in order to bypass G-quadruplex structures that block replisomes (Koole et al., 2014; Van Schendel et al., 2015). G-quadruplexes that form during mammalian telomere replication result in fragile telomeres that are apparent in metaphase chromosome spreads and occur more frequently when RTEL1 is deficient (Vannier et al., 2012). If DNA polymerase theta normally creates small deletions that bypass G-quadruplexes during mammalian telomere replication, then this promiscuous DNA polymerase may be specially licensed to operate at dysfunctional telomeres. If so, DNA polymerase theta might be naturally suited to responding to short telomeres that are unable to form a T-loop.

Perfect telomere sequences within ITS tracts that were 9-24 nucleotides in length were often targeted for DNA synthesis by short telomeres. However, we identified two telomere-ITS transitions that occurred at short 4 nucleotide stretches of telomere homology, which might be consistent with the short primer requirements of DNA polymerase theta. In addition, we identified a 6 nucleotide insertion and two point mutations at three telomere-ITS DNA synthesis transitions (Fig. 2D). The latter three mutations are consistent with template slippage events that occur when DNA polymerase theta acts to create small insertions and deletions that are frequently observed at TMEJ breakpoints (Wood & Doublié, 2016). We also observed small insertions characteristic of DNA polymerase theta at the borders of template shifting events that occurred at later stages of subtelomeric DNA synthesis (Fig. 2E). Together, these data imply that DNA polymerase theta acts during initial and subsequent subtelomeric DNA synthesis events that are separated by bouts of template shifting, in addition to sealing the subtelomeric duplication with a distinct chromosome end (Fig. 5B). Consistently, subtelomeric template shifting events that lacked insertions possessed 1 or 2 nucleotides of homology characteristic of DNA polymerase theta at their borders, and mutation of *polq-1* strongly repressed both telomere-ITS DNA synthesis and telomere fusion.

We studied telomere fusions that occurred in the genomes of *C. elegans* ALT strains (E. Kim et al., 2021) and identified independent examples of template shifting that occurred in the context of subtelomeric DNA synthesis. In one case, 16 nucleotides from an ITS tract were added to a short telomere, and then a template-shift resulted in DNA synthesis at subtelomere sequence based on 2 nucleotides of homology (Fig. 2F). In another case, a subtelomeric DNA synthesis event at IIR transitions to the center of Chromosome II based on 2 nucleotides of homology and copies 15 nucleotides of DNA that are used to seal the telomere fusion with VL (Fig. 2G). These events are consistent with the ability of DNA polymerase theta to create insertions in the context of end-joining by copying short stretches of DNA from a linked locus or a different chromosome.

Given that both *trt-1 rtel-1* mutants and *trt-1; polq-1* mutants lack chromosome fusions and display strong reductions in telomere-ITS DNA synthesis, we asked if RTEL-1 might promote TMEJ of CRISPR/Cas9-induced DSBs, which are known to depend on POLQ-1 (Van Schendel et al., 2015). We found that POLQ-1 was more efficient at generating CRISPR/Cas9-induced deletions when *rtel-1* was mutant, which is consistent with known roles of RTEL-1 as an anti-recombinase (Barber et al., 2008; León-Ortiz et al., 2018). We therefore conclude that POLQ-1 and RTEL-1 are likely to play distinct roles in orchestrating subtelomeric DNA synthesis when telomeres are too short to create a T-loop. DNA polymerase theta appears to be responsible for initiation of DNA synthesis, at least in some cases. DNA polymerase delta could continue DNA synthesis based on the primer created by DNA polymerase theta. RTEL-1 would then dismantle this DNA synthesis intermediate and hand the 3’ overhang to DNA polymerase theta, which could continue subtelomeric DNA synthesis (by initiating a template shift) or could create a chromosome-type telomere fusion by sealing the subtelomeric duplication with the subtelomere of a distinct chromosome (Fig. 5B).

We found that loss of either POLQ-1 or RTEL-1 repressed the creation of telomere fusions that normally occur after 15 to 20 generations of growth of *trt-1* telomerase mutants, and telomere fusions were not even observed after 100 generations of growth of *trt-1 rtel-1* double mutant strains that maintain their telomeres by ALT. These data imply that RTEL-1/RTEL1 and DNA polymerase theta play essential roles in the creation of most telomere fusions that occur in the absence of telomerase, at least in part by orchestrating sub-telomeric DNA synthesis events.

Degenerate ITS tracts are commonly found scattered on metazoan chromosome arms. Although the origin of ITS tracts remains a mystery (Frenk et al., 2019), telomere-ITS DNA synthesis events have been found to initiate ALT in *C. elegans* telomerase mutants, resulting in telomeres that alternate between short tracts of telomeric DNA, ITS tract DNA, and unique sequence DNA that is next to an ITS tract (E. Kim et al., 2021; Seo et al., 2015). Our study suggests that a second important function of ITS tracts is to promote the creation of telomere fusions that occur in the absence of telomerase. Subtelomeric DNA synthesis events are therefore likely to play prominent roles in shaping human tumor genomes. On an evolutionary time-scale, stochastic subtelomeric DNA synthesis events that are initiated by telomeres may help to explain the repetitive and rapidly evolving nature of sub-telomeric DNA (C. Kim et al., 2019).

## Materials and Methods

### Strains

Worms were propagated at 20°C on Nematode Growth Medium (NGM) plates seeded with OP50 *E. coli*. Strains used include: Bristol N2 wild type, YA1059 *trt-1(ok410) I*, TG2228 *polq-1(tm2026) III*, TG1663 *ercc-1(tm1981) I*, *rtel-1(tm1866) I, rtel-1(yp15) I*, *rtel-1(yp16) I*, *rtel-1(yp17) I*, *rtel-1(yp18) I*, *ercc-1(tm1981) I*, *ercc-1(tm2013) I*, and strains containing ypT29 and ypT50 fusions (Lowden, 2011). *trt-1* mutants display a mortal germline phenotype, so passaging for maintenance was accomplished using either an *unc-29* genetic marker or *hT2* balancer.

### Plasmids

pELS16 [Peft-3::Cas9 + dpy-10 sgRNA], pELS50 [Peft-3::Cas9 + rtel-1 sgRNA], pELS51 [Peft-3::Cas9 + rtel-1 sgRNA], pELS52 [Peft-3::Cas9 + rtel-1 sgRNA], and pELS53 [Peft-3::Cas9 + rtel-1 sgRNA] plasmids were generated for this study by modifying the pDD162 plasmid (Dickinson, 2013) using Q5 Site Directed Mutagenesis Kit (NEB, Cat# E0554S, Ipswich, MA).

### Generation of new strains by CRISPR

New *rtel-1* mutant alleles were generated in a *trt-1(ok410)* background using CRISPR based on previously described protocols (Dickinson et al., 2013). Injected lines were screened with the help of *dpy-10* co-CRISPR, based on previously described protocols (Arribere et al., 2014). Identified mutations were characterized by Sanger sequencing (Fig. S4A). The *yp15* allele includes a premature stop codon downstream of the DEAH box that is predicted to eliminate 60% of the *rtel-1* coding region. The *yp18* allele contains a 228 amino acid deletion that abolishes the DEAH box. Both of these predicted loss-of-function alleles show strong reductions in progeny production. *yp16* and *yp17* alleles are small in-frame indels which are expected to encode functional *rtel-1*.All alleles were crossed as soon as possible to males containing the *hT2* balancer, which balances the *trt-1* and *rtel-1* loci and allows for the stable propagation of heterozygotes.

Progeny production was measured in all CRISPR-based *rtel-1* mutant strains to check for phenocopying of the *rtel-1(tm1866)* allele, in which progeny production is dramatically reduced (Fig. S4B). Progeny production was similarly reduced for *yp15* and *yp18* alleles. The expected function *yp16* and *yp17* alleles had significantly higher progeny production, although fertility data indicates that *yp16* is a hypomorphic allele.

### Germline mortality assay

Germline mortality was assayed using previously described procedures (Cheng et al., 2012). Heterozygous telomerase mutants were maintained using either an *unc-29* marker or *hT2* balancer. Mortality assays began upon homozygousing the mutant telomerase allele. Individual lines were established by putting 6 L1 larvae onto plates. 6 L1 larvae were then passaged weekly until sterility.

ALT assays were carried out similarly, except that rather than individual worms a 1-cm square of agar was cut out and transferred to new plates weekly. These agar squares can contain hundreds of worms. Sterility was considered reached when fewer than 6 larvae could be found on a plate.

### ITS-DNA synthesis assay

PCR for the ITS-DNA synthesis assay was carried out using ExTaq polymerase (Takara Bio cat# RR001A, Kasatsu, Japan). 35 cycles were executed with 64°C annealing temperature and 1 minute extension time. Primer sets were composed of one “ITS” and one “Telomere” primer (Table S3). Samples were run on 1% agarose gels containing ethidium bromide.

### Whole genome sequencing and analysis

Purified DNA was subjected to 50bp paired-end sequencing on an Illumina HiSeq 4000. Reads were adaptor-trimmed using bbduk.sh from the bbmap package (http://sourceforge.net/projects/bbmap) and aligned to the C. elegans genome (WS251) using BWA-mem (https://github.com/lh3/bwa). Structural variant calling was performed using SvABA (https://github.com/walaj/svaba) and copy number changes were detected using CNVnator (http://sv.gersteinlab.org/cnvnator). Genomic regions were visualized using the Gviz R package (https://github.com/ivanek/Gviz), BBMap short-read aligner, and other bioinformatics tools.

### Southern blot

Southern blotting was performed based on previously published protocols (Ahmed, 2000). Probe was generated using the PCR DIG probe synthesis reaction kit (Roche cat# 11636090910, Basel, Switzerland). Tel2 (GAATAATGAGAATTTTCAGGC) and T3 (CGAAATTAACCCTCACTAAAG) primers were used to amplify off of cTel55x plasmid template (Wicky, 1996).

### Statistics

P-values for ITS-DNA synthesis and progeny production assays were computed using the Wilcox test. P-values for germline mortality data were calculated using the log-rank test. P-values for chromosome number were computed using Welch’s t-test. Multiple hypothesis adjustment was made using the Holm method (Holm, 1979). Unless otherwise noted, all error bars represent 95% confidence intervals. Significance values were computed using R.

### Data availability

High throughput sequencing data sets are posted on the NCBI Sequence Read Archive (SRA) database, and are publicly accessible at https://www.ncbi.nlm.nih.gov/sra/PRJNA839276.

## Author contributions

E.L., M.B. B.H., S.F., A.M., M.N., A.-M.L.O. and S.A. performed experiments. E.L., M.B. S.F. and B.H. analyzed the data. S.F., E.L., and S.A. wrote the manuscript with editorial input from S.J.B.

## Acknowledgements

S.A. is supported by NIH R01 GM135470. A.-M.L.O. and S.J.B. are supported by the Francis Crick Institute, which receives its core funding from Cancer Research UK (FC0010048), the UK Medical Research Council (FC0010048) and the Wellcome Trust (FC0010048). S.J.B is also funded by a European Research Council (ERC) Advanced Investigator Grant (TelMetab); and Wellcome Trust Senior Investigator and Collaborative Grants.

---

**Fig. S1. Related to Figure 1.**
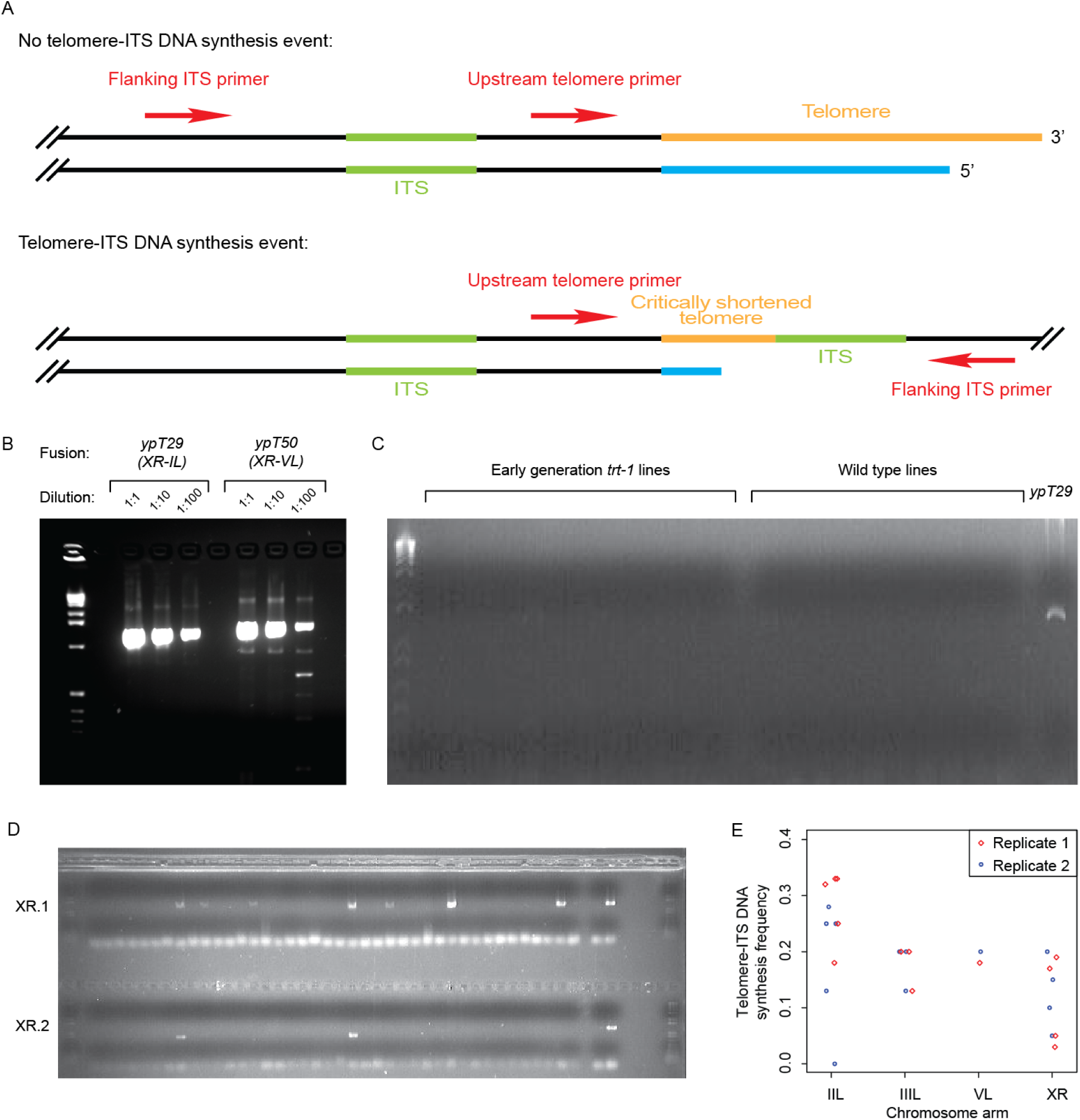
(A) Diagram of general primer positions for the telomere-ITS DNA synthesis assay. (B) The telomere-ITS DNA synthesis assay was able to detect chromosome fusions in worm lines stably propagating fused chromosomes. Detection was robust even when DNA from the fusion lines was diluted 1:100 with wild-type DNA. (C) Fusions are not detected in wild type or early generation *trt-1* mutants, but are detected in lines containing chromosome fusions. (D) Telomere-ITS DNA synthesis events are detected in late generation *trt-1* mutants. (E) Telomere-ITS DNA synthesis frequency was not significantly different between two sets of 40 *trt-1* mutant lines (p=0.24 by paired t-test).

**Fig. S2.**
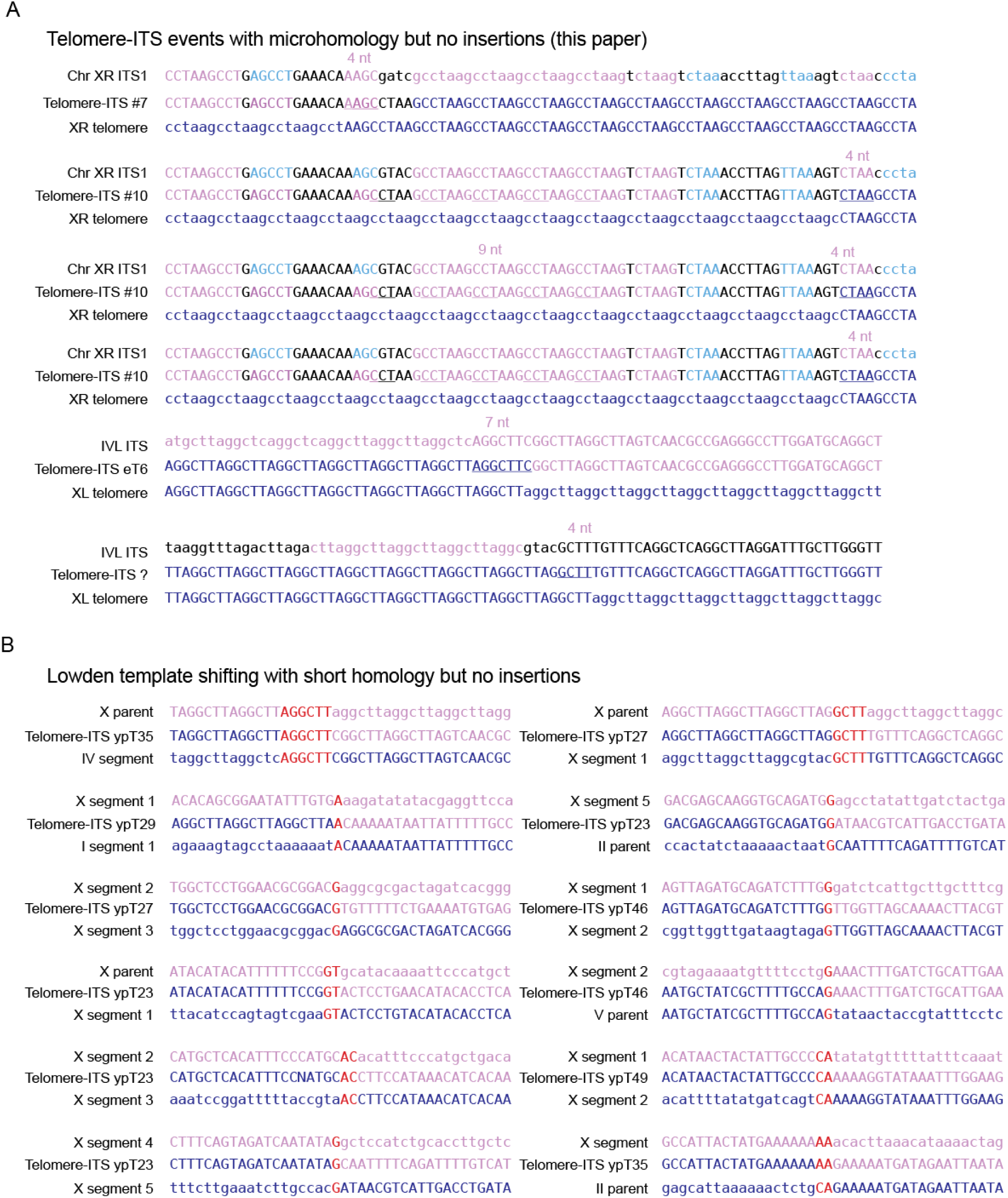
Microhomologies at subtelomeric DNA synthesis transitions. Microhomology present at (A) telomere-ITS DNA synthesis events and (B) template-shifting events, based on previously published sequences (Lowden et al., 2011).

**Fig. S3.**
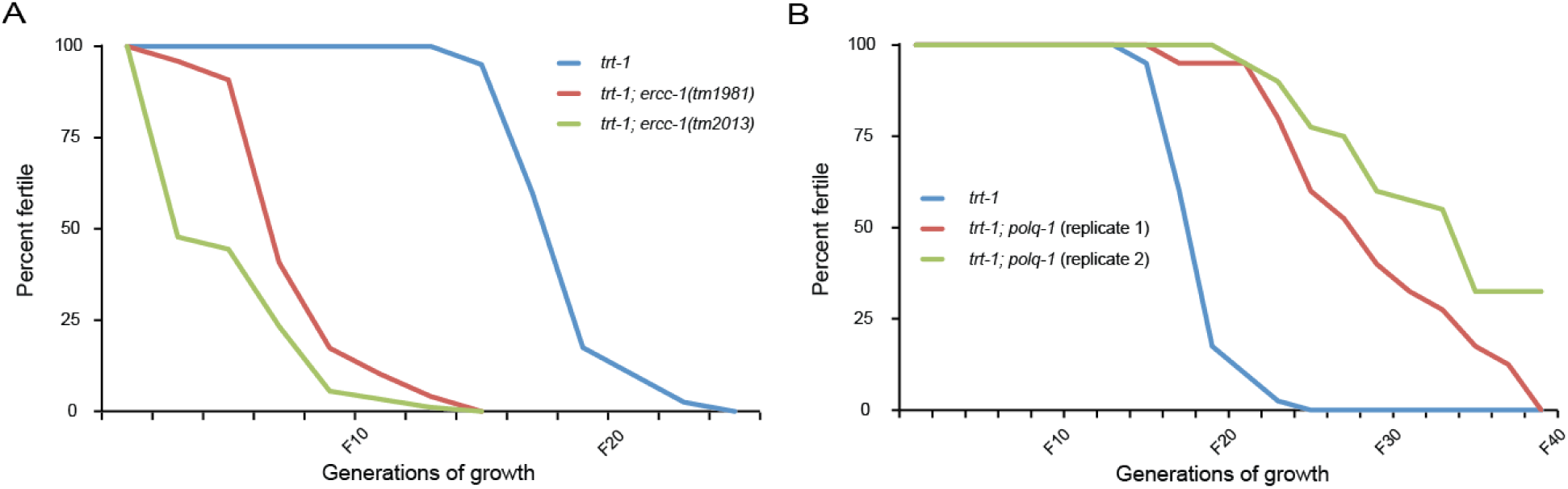
Transgenerational mortality phenotypes for double mutants. (A) *trt-1; ercc-1* double mutant lines went sterile more rapidly than *trt-1* single mutants. (B) *trt-1; polq-1* double mutant lines took longer to go sterile than *trt-1* single mutants.

**Fig. S4.**
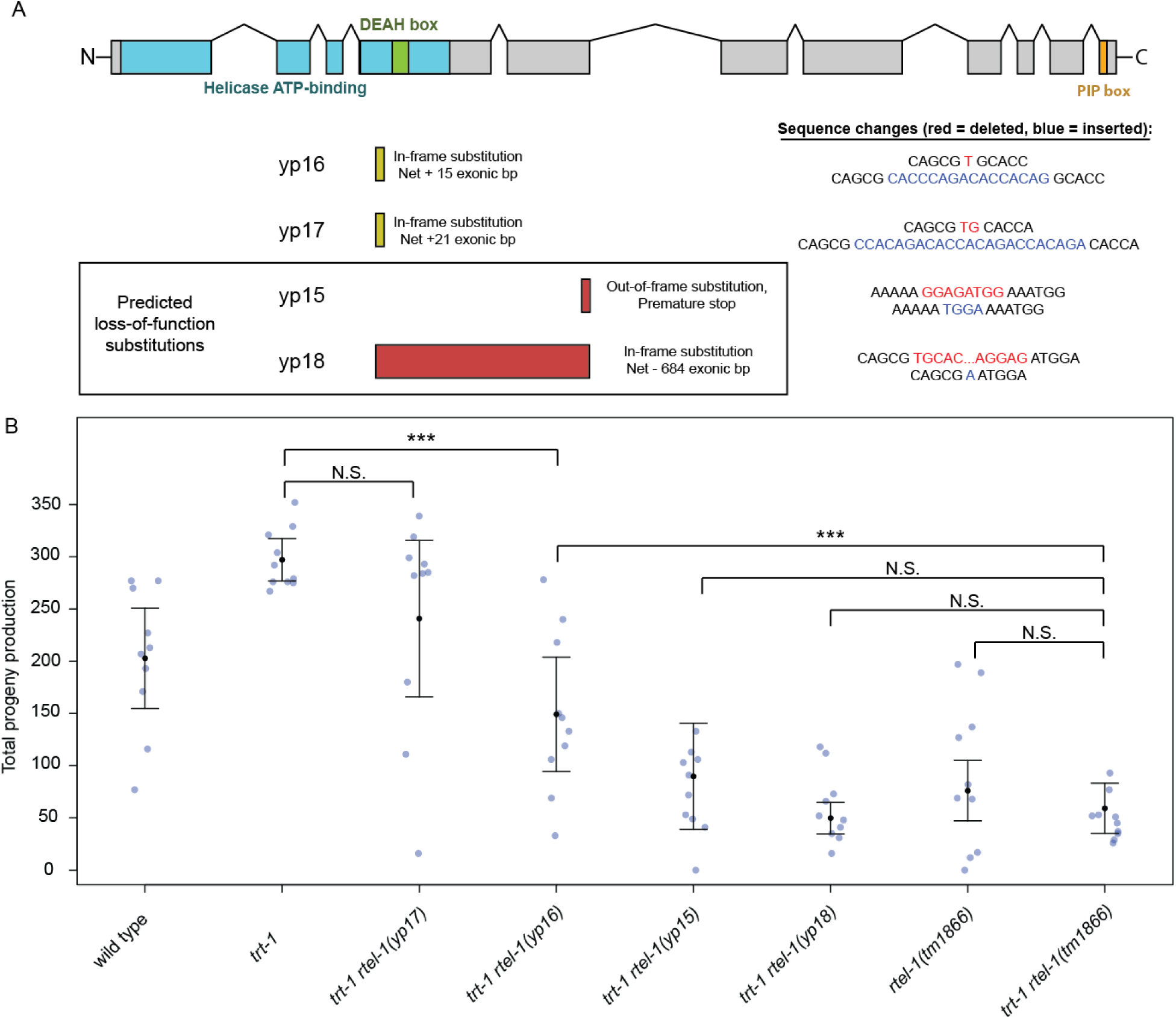
(A) Diagram of mutant *rtel-1* alleles generated by CRISPR. (B) Progeny production in strains possessing CRISPR-based *rtel-1* mutant alleles as well as the previously described allele *tm1866*. The frame-shift alleles *yp15* and *yp18* reduced progeny production to a similar extent as *tm1866* in a *trt-1* mutant background. Indicated significance is in comparison to *trt-1 rtel-1(tm1866)* single mutants. ** p < 0.01, *** p < 0.005.

## Supplemental Information

### Additional Files

Figure 1-figure supplement 1B-source data 1

Figure 1-figure supplement 1B-source data 1-data supplement

Figure 1-figure supplement 1C-source data 1

Figure 1-figure supplement 1C-source data 1-data supplement

Figure 1-figure supplement 1D-source data 1

Figure 1-figure supplement 1D-source data 1-data supplement

Figure 3-source data 1

Figure 3-source data 1-data supplement

**Table S1:**
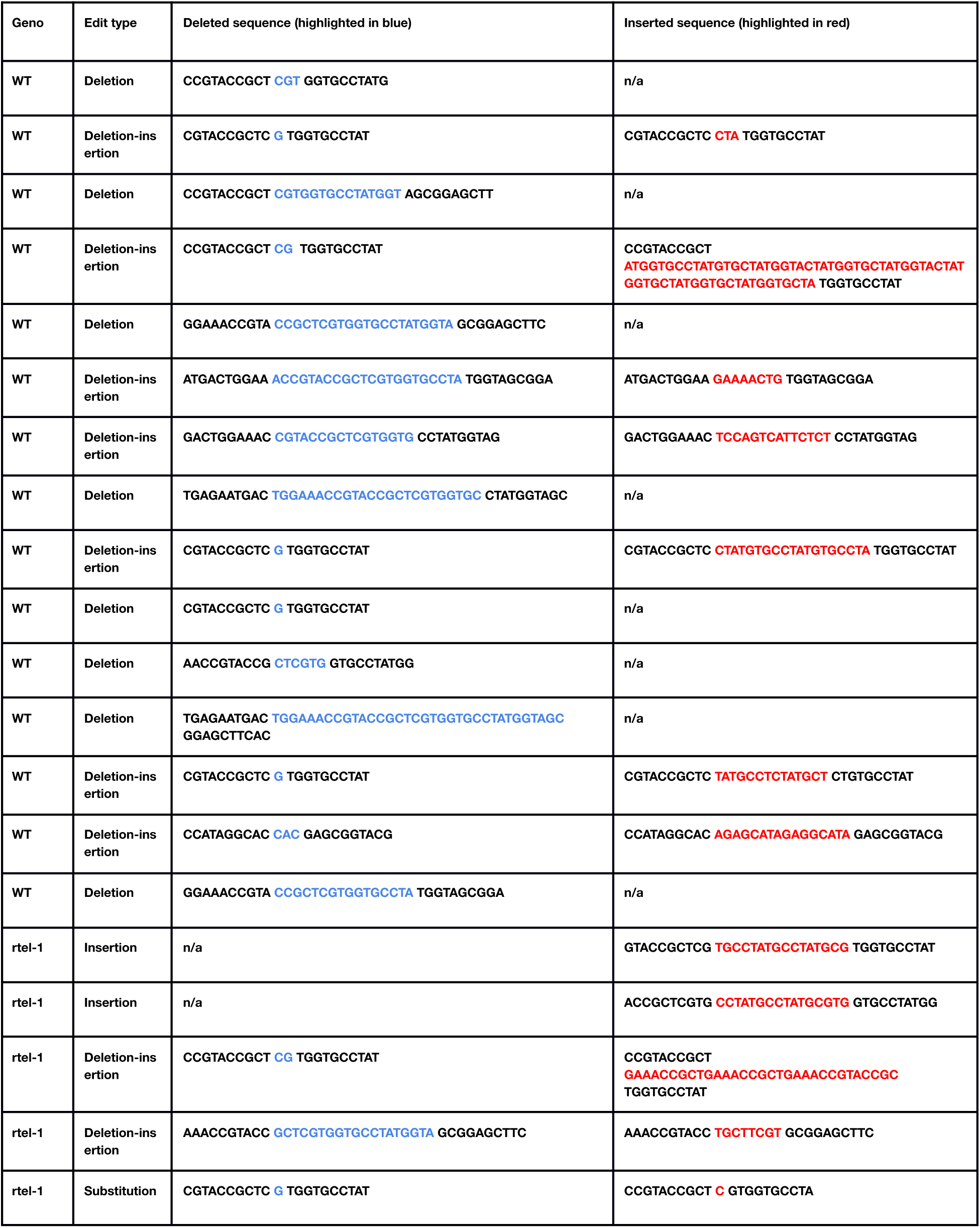
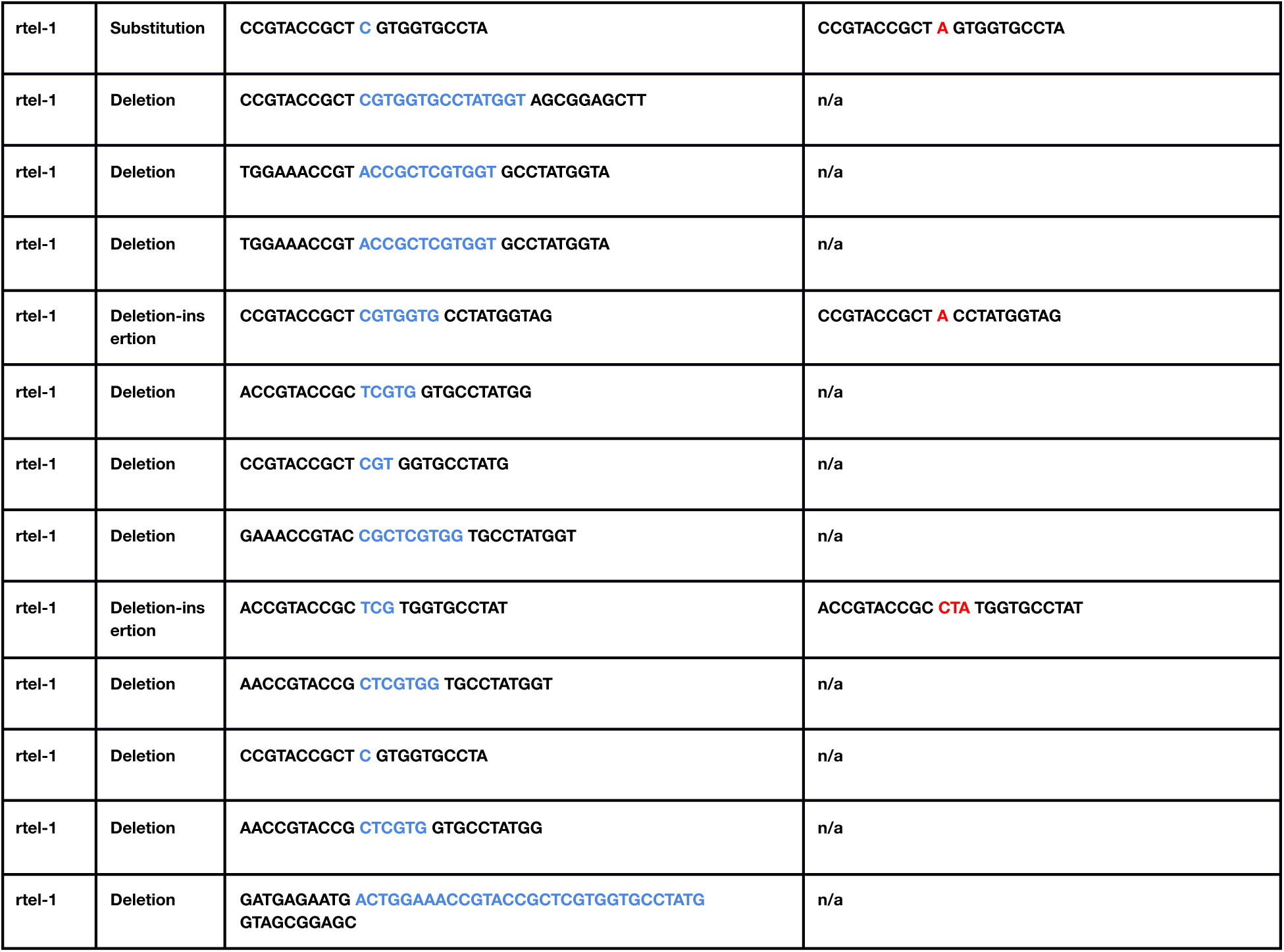
Sequence changes for CRISPR edits of *dpy-10* loci in wild-type and *rtel-1* mutant worms.

**Table S2:** Positions of ITSs that border genome copy number changes based on whole genome sequence analysis of telomerase mutants.

| chrom | start | end | width | strand | count | score | n_events | pcr | pcr_recomb |
| --- | --- | --- | --- | --- | --- | --- | --- | --- | --- |
| I | 577 | 846 | 270 | - | 28 | 91.1 | 1 | No | 0 |
| I | 13903202 | 13903332 | 131 | - | 10 | 87.9 | 1 | No | 0 |
| I | 14440692 | 14441148 | 457 | + | 26 | 73.8 | 3 | No | 0 |
| II | 4013236 | 4013502 | 267 | + | 7 | 70.6 | 1 | No | 0 |
| II | 14117029 | 14117282 | 254 | - | 14 | 67.4 | 1 | No | 0 |
| II | 15204163 | 15204348 | 186 | - | 10 | 84.4 | 1 | No | 0 |
| IV | 1103302 | 1103381 | 80 | - | 4 | 76.5 | 2 | No | 0 |
| IV | 3590527 | 3590692 | 166 | - | 4 | 62.7 | 2 | No | 0 |
| IV | 16172768 | 16173145 | 378 | - | 20 | 83.4 | 1 | No | 0 |
| V | 16075 | 16349 | 275 | + | 8 | 76.6 | 1 | Yes | 18 |
| V | 248722 | 248877 | 156 | - | 8 | 74.1 | 1 | No | 0 |
| V | 365708 | 365827 | 120 | + | 8 | 86.7 | 4 | No | 0 |
| V | 1122552 | 1122857 | 306 | - | 15 | 83.3 | 1 | No | 0 |
| V | 1134184 | 1134389 | 206 | - | 13 | 85.3 | 1 | No | 0 |
| V | 1142352 | 1142532 | 181 | - | 11 | 84.5 | 1 | No | 0 |
| V | 1423358 | 1423759 | 402 | - | 37 | 91 | 1 | No | 0 |
| V | 1480406 | 1480585 | 180 | - | 15 | 87.8 | 1 | No | 0 |
| V | 1481685 | 1481924 | 240 | - | 15 | 83.8 | 1 | No | 0 |
| V | 1826256 | 1826420 | 165 | + | 4 | 48.3 | 1 | No | 0 |
| V | 4684246 | 4684415 | 170 | - | 14 | 82.4 | 2 | No | 0 |
| V | 17552594 | 17552707 | 114 | - | 8 | 84.2 | 1 | No | 0 |
| V | 19517962 | 19518331 | 370 | + | 29 | 86.9 | 1 | No | 0 |
| V | 19996804 | 19997352 | 549 | - | 21 | 75 | 2 | No | 0 |
| X | 17578021 | 17578200 | 180 | + | 10 | 82.8 | 2 | Yes | 0 |
| X | 17632898 | 17632933 | 36 | + | 4 | 91.7 | 1 | No | 0 |

**Table S3:**
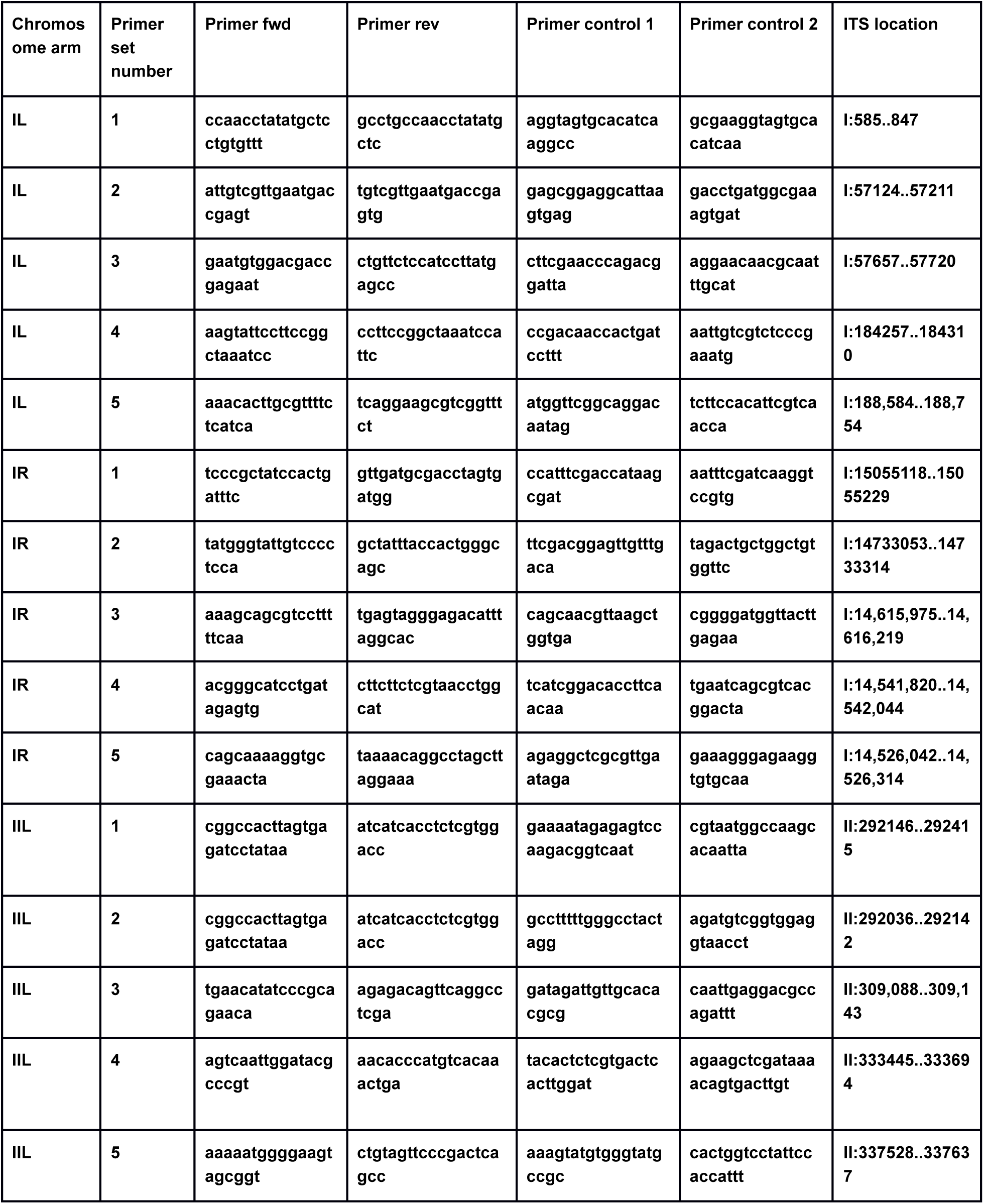
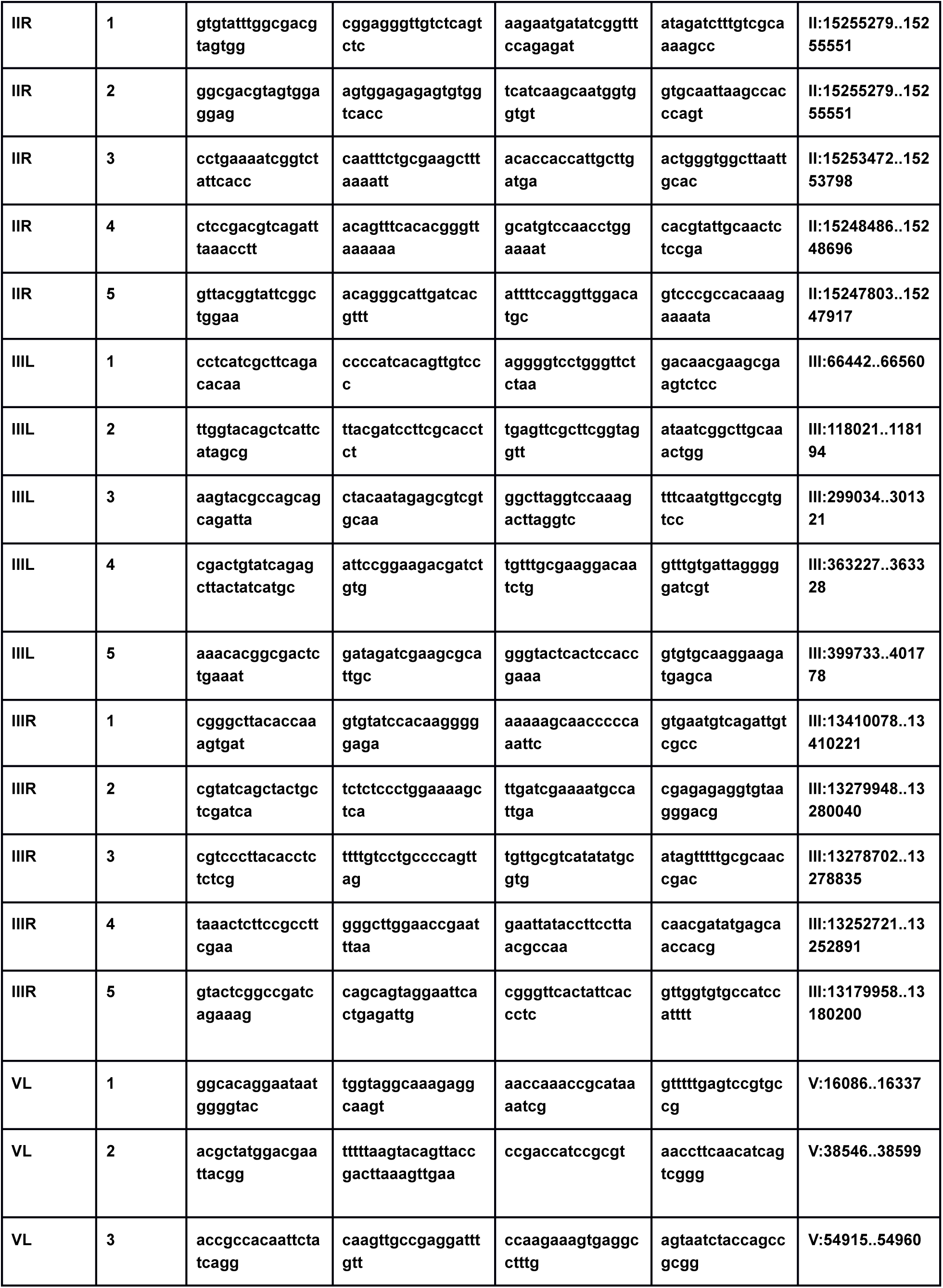
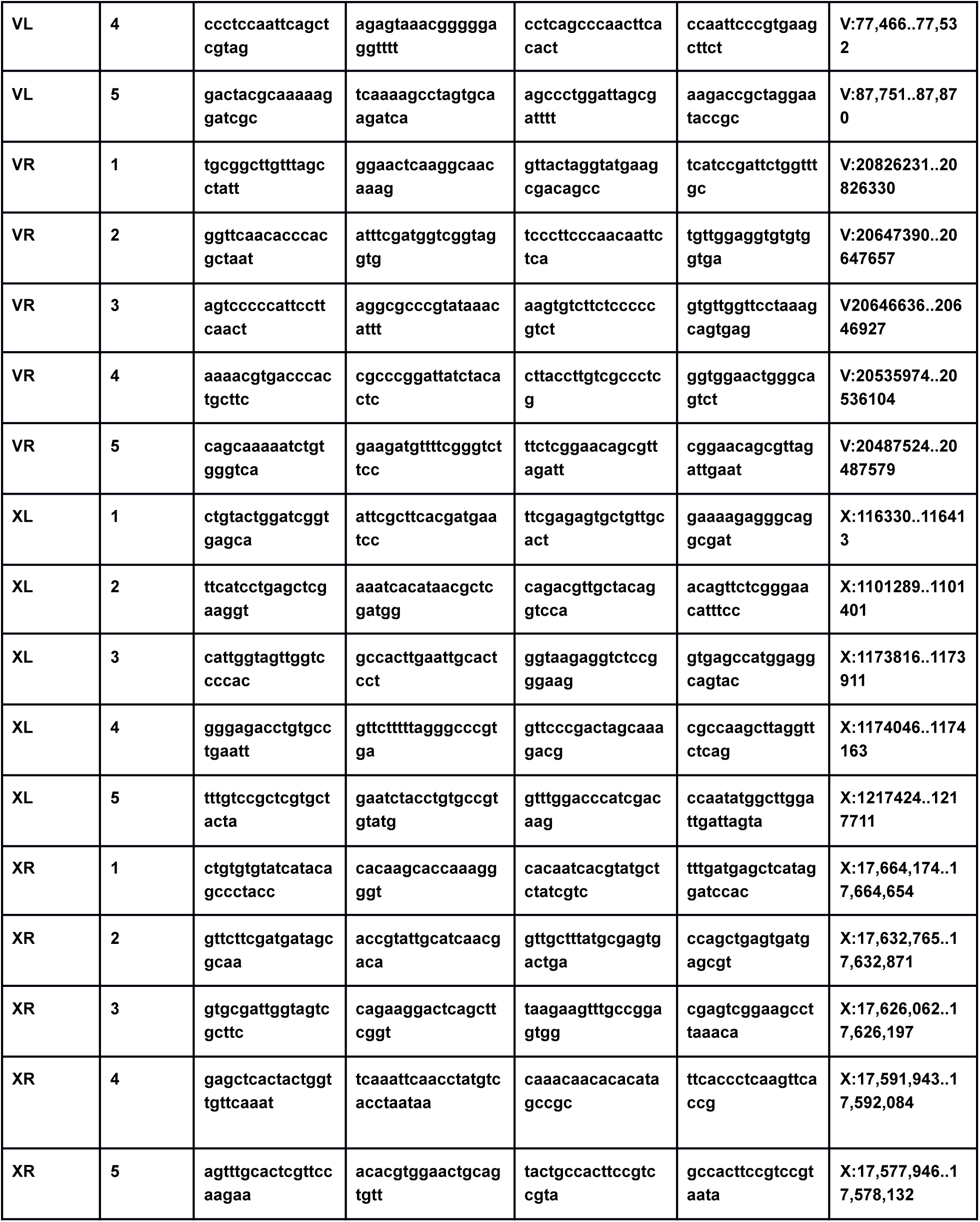
Primers used for PCR-based telomere-ITS DNA synthesis assay. Forward primers were used in combination with primers facing the telomere. Forward and Reverse ITS primers that flank each ITS are indicated, which were combined to ensure that robust PCR products were obtained under conditions where telomere-ITS DNA synthesis events were tested.

